# Sequential accumulation of dynein and its regulatory proteins at the spindle region in the *Caenorhabditis elegans* embryo

**DOI:** 10.1101/2021.09.14.460385

**Authors:** Takayuki Torisawa, Akatsuki Kimura

## Abstract

Cytoplasmic dynein is responsible for various cellular processes during the cell cycle. The mechanism by which its activity is regulated spatially and temporarily inside the cell remains elusive. There are various regulatory proteins of dynein, including dynactin, NDEL1/NUD-2, and LIS1. Characterizing the spatiotemporal localization of regulatory proteins *in vivo* will aid understanding of the cellular regulation of dynein. Here, we focused on spindle formation in the *Caenorhabditis elegans* early embryo, wherein dynein and its regulatory proteins translocated from the cytoplasm to the spindle region upon nuclear envelope breakdown (NEBD). We found that (i) a limited set of dynein regulatory proteins accumulated in the spindle region, (ii) the spatial localization patterns were distinct among the regulators, and (iii) the regulatory proteins did not accumulate in the spindle region simultaneously but sequentially. Furthermore, the accumulation of NUD-2 was unique among the regulators. NUD-2 started to accumulate before NEBD (pre-NEBD accumulation), and exhibited the highest enrichment compared to the cytoplasmic concentration. Using a protein injection approach, we revealed that the C-terminal helix of NUD-2 was responsible for pre-NEBD accumulation. These findings suggest a fine temporal control of the subcellular localization of regulatory proteins.

## INTRODUCTION

Cytoplasmic dynein I is a molecular motor that moves along a microtubule towards its minus-end (King, 2011). Cytoplasmic dynein I is responsible for and plays a vital role in a substantial extent of the minus-end-directed transport in animal cells (Roberts *et al.*, 2013). In this study, we referred to cytoplasmic dynein I as dynein for simplicity. A striking feature of dynein is that the heavy chain, including ATP hydrolysis sites and a microtubule-binding domain, is encoded by a single gene (Pfister *et al.*, 2006). This is in contrast to kinesins, which are also recognized as microtubule-based motors that mostly move towards the plus-end of microtubules. In case of kinesins, there are multiple genes encoding kinesin motors that share a similar motor domain; additionally, specific types of kinesin are expressed at specific times and are localized to specific regions to perform specific functions (Hirokawa *et al.*, 2009; Cross and McAinsh, 2014). To demonstrate dynein-associated cellular functions, it is imperative that the cell utilizes a single type of dynein heavy chain at various times and locations in a regulated manner. Therefore, the regulation of the localization, timing, and activity of dynein should be sophisticated and should occur at multiple levels, from intramolecular regulation to regulation at the population level (Kardon and Vale, 2009; Reck-Peterson *et al.*, 2018; Torisawa and Kimura, 2020). An example of intramolecular regulation is an autoinhibition mechanism in which the isolated, solely existing dynein tends to assume a characteristic phi-shaped form and only shows diffusive movements along microtubules (Torisawa *et al.*, 2014; Zhang *et al.*, 2017).

Dynein is associated with various regulatory proteins. Dynactin is a major regulatory protein for dynein. Dynactin associates with dynein and demonstrates the formation of a complex at various cellular sites (Reck-Peterson *et al.*, 2018). The formation of a complex with dynactin is one of the mechanisms to release dynein from the autoinhibited state, and it provides the basis for the formation of larger complexes with various regulatory proteins (Olenick and Holzbaur, 2019). In addition to dynactin, several regulatory proteins, such as LIS1, NDEL1/NDE1, and NuMA exist. LIS1 controls the force generation of dynein and aids formation of the dynein-dynactin complex (McKenney *et al.*, 2010; Elshenawy *et al.*, 2020; Htet *et al.*, 2020; Marzo *et al.*, 2020). NDE1/NDEL1 is known to aid the establishment of interaction between LIS1 and dynein (Yamada *et al.*, 2008; McKenney *et al.*, 2010; Torisawa *et al.*, 2011). NuMA catalyzes the recruitment of dynein to the cell cortex to facilitate the formation of cortical force generators (Kiyomitsu, 2019). The dynein molecules for the complex should be reserved at the bulk cytoplasm; however, the mechanism and the timing for the formation of the complex remain elusive. In many organisms, dynein is found throughout the cytoplasm (Portegijs *et al.*, 2016; Schmidt *et al.*, 2017; Heppert *et al.*, 2018). However, a pertinent question exists: do such dyneins exhibit the formation of a complex with regulatory proteins before being recruited to specific functional sites? Such a question is especially important when a new microtubule-associated structure is formed in a cell according to temporal cues.

A prominent example of the temporal and spatial regulation of dynein is the formation of mitotic spindles. Dynein is excluded from the nucleus during interphase, but is incorporated into the spindle during mitosis and plays an important role in spindle formation and function (Vaisberg *et al.*, 1993). In spindle formation, nuclear envelope breakdown (NEBD) enables the movement of cytoplasmic molecules into the nuclear region. Dynein and the regulatory proteins involved in spindle formation, such as dynactin, NuMA, LIS1, and NDEL1/NDE1, are translocated into the region (Raaijmakers *et al.*, 2013). Apart from the proteins related to dynein, spindle component proteins, including tubulin and associated proteins, also present with accumulation in the spindle region. In the *Caenorhabditis elegans embryo*, the dynein heavy chain, as well as tubulin and other molecules, were observed to undergo accumulation in the nuclear area after NEBD, but such events were independent of spindle formation (Hayashi *et al.*, 2012). We proposed that tubulin and other molecules could accumulate in the area before spindle formation and referred to this transiently formed area as ‘nascent spindle region’ (Hayashi *et al.*, 2012). The detailed timing and localization of the proteins undergoing accumulation in the nascent spindle region have not been examined thus far. It has been naively assumed that these proteins translocate to the nascent spindle region simultaneously upon NEBD without specific regulations.

In this study, we analyzed the accumulation of dynein and its regulatory proteins at the spindle region to understand the mechanism by which spatiotemporal regulation of dynein was achieved during spindle formation in *Caenorhabditis elegans* embryos. Quantitative analysis of the accumulation phenomena showed variations in the initial events of accumulation, the maximum accumulated amount, and the accumulation rate. Chemical perturbation revealed that the proteins also differed in the accumulation sites within the spindle region, including the spindle microtubules, chromosomes, and/or bulk nucleoplasm. Among the proteins analyzed, NUD-2, a *C. elegans* ortholog of NDEL1/NDE1, showed a characteristic accumulation that commenced before NEBD. This earlier accumulation process was observed to be dependent on the Ran GTPase activity. The depletion of NUD-2 reduced the ability of the spindle region to retain the accumulated proteins, but it did not affect the accumulation process itself. Furthermore, using the injection technique for the recombinant proteins, we found that the C-terminal helix region of NUD-2 was necessary for its accumulation before NEBD. Our results suggest the implication of the accumulation phenomena for the spatiotemporal regulation of cytoplasmic dynein during the formation of mitotic spindles.

## RESULTS

### Accumulation of endogenously tagged dynein and its regulators during spindle formation in *C. elegans* early embryos

To investigate the spatiotemporal regulation of dynein and its regulatory proteins during spindle formation, we observed accumulation events of dynein and its regulatory proteins in *C. elegans* early embryos. Previous studies have shown that dynein and certain regulatory proteins are localized in the spindle region, as evidenced via transgene expression (Cockell *et al.*, 2004; Hayashi *et al.*, 2012). In this study, we used specific worm strains expressing the heavy chain of cytoplasmic dynein I, the p150 subunit of dynactin complex, and several other regulatory proteins, including LIS-1, NUD- 2/NDEL1, and LIN-5/NuMA, from the endogenous locus (**Table S1**). These proteins were selected because they have been known to be involved in mitosis, with presence reported in the cytoplasm during interphase. Other regulatory proteins, including SPDL-1/Spindly and NUD-1/NudC, are primarily localized in the nucleus (Aumais *et al.*, 2003; Gassmann *et al.*, 2008). We constructed a new worm strain expressing hsGFP-tagged DHC-1 using the CRISPR/Cas9 method (see Materials and Methods). To observe other proteins, we utilized the strains reported in previous studies (Heppert *et al.*, 2018).

Using confocal microscopy, we observed the NEBD-dependent accumulation of dynein, dynactin, LIN-5, LIS-1, and NUD-2 in the spindle region (**Figure 1A; Movie S1**). In *C. elegans*, nuclear pore complexes (NPCs) undergo disassembly that is initiated in prophase, and the permeability barrier between the nucleoplasm and cytoplasm is lost in prometaphase (Lee *et al.*, 2000; Tzur and Gruenbaum, 2013), thereby enabling diffusion of the mitotic proteins in the cytoplasm into the spindle region. Based on the permeability, we referred to this change in the nucleocytoplasmic barrier as *C. elegans* nuclear envelope breakdown (CeNEBD) in a previous study (Hayashi *et al.*, 2012). To establish controls, we considered BICD-1 and ZYG-12 as candidates for non-accumulating proteins. BICD-1 is known as a single *C. elegans* ortholog of the BicD family protein (Aguirre-Chen *et al.*, 2011), and ZYG-12 is considered as the *C. elegans* ortholog of the Hook family proteins (Malone *et al.*, 2003). Both BicD and Hook are recognized as adaptor proteins of dynein for intracellular transport (Olenick and Holzbaur, 2019) and have been hypothesized to play minor roles in mitosis. As expected, we did not observe accumulation of BICD-1 or ZYG-12. A signal of BICD-1 was not detected as previously described (Heppert *et al.*, 2018); we only observed the autofluorescence signals in the GFP channel (Heppert *et al.*, 2016). In contrast, we found that ZYG-12 existed in the early embryos and was localized mainly in the nuclear membrane, as previously described (Malone *et al.*, 2003).

**Figure 1.**
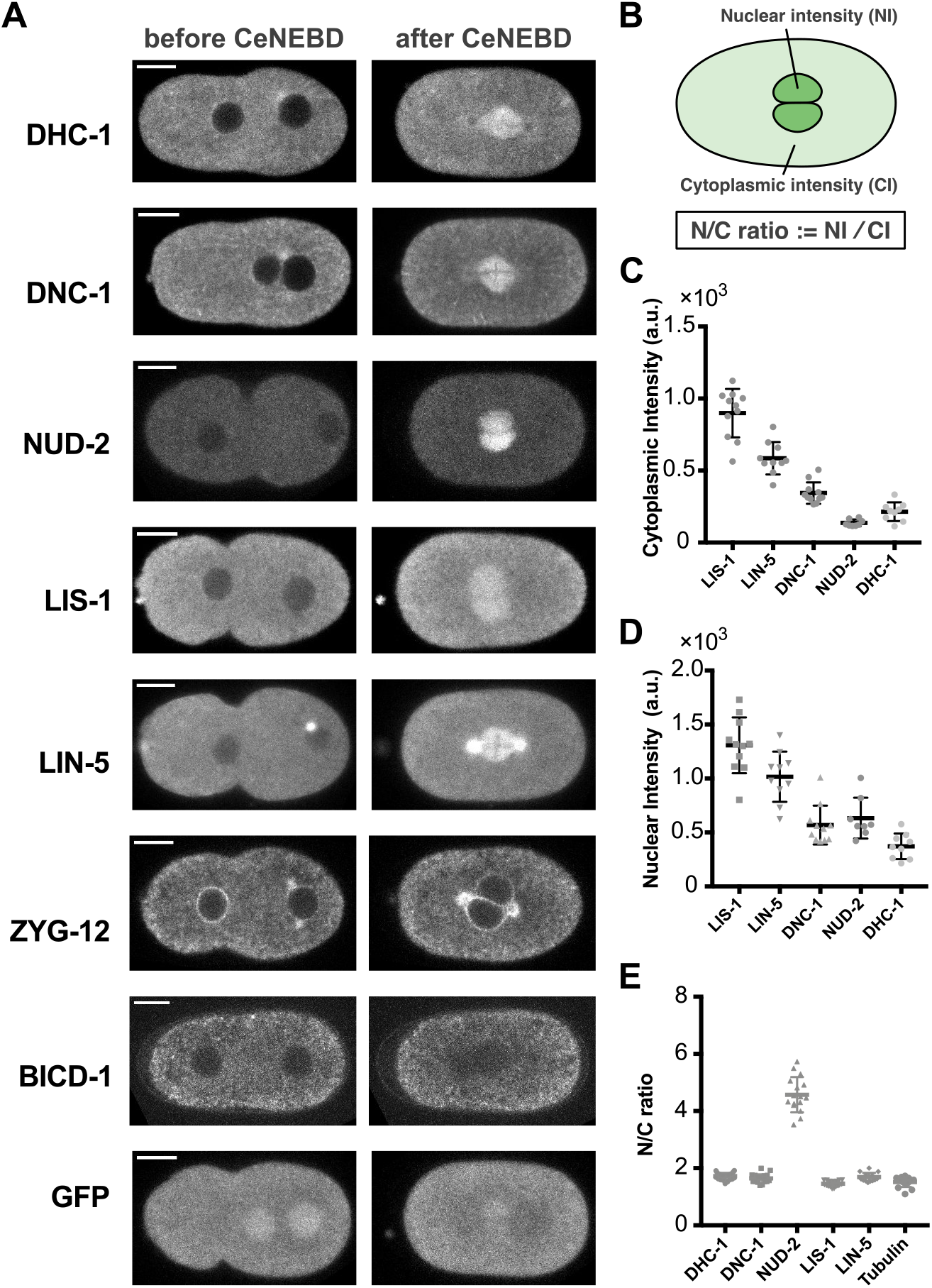
Observations of the temporal dynamics of dynein and its regulatory proteins during the 1st mitosis of *Caenorhabditis elegans* early embryos. (A) Typical single-plane time-lapse images showing the cellular localization of dynein and its regulatory proteins before and after CeNEBD. Scale bars, 10 μm. (B) Schematic representation of quantification of accumulation phenomenon. Nuclear intensity (NI) and Cytoplasmic intensity (CI) indicate the mean intensity of the nuclear region and cytoplasmic region, respectively. (C-E) Quantification of the accumulated amount of dynein, dynactin, LIS-1, NUD-2, and LIN-5. (C) The mean cytoplasmic intensity. (D) The maximum nuclear (spindle) intensity. (E) The maximum N/C ratio (i.e., the intensity in the nuclear/spindle region divided by the cytoplasmic intensity.

Observations of dynein and regulatory proteins expressed from endogenous loci enabled the quantification of the amounts and stoichiometry of the accumulated proteins. In the analysis, we assumed that the expression levels of the endogenously tagged proteins were not largely affected by the tags, and that the amount of protein and the fluorescence intensity exhibited a linear relationship. We first quantified the signal intensity in the cytoplasm before pronuclear formation (cytoplasm intensity or CI) as an index reflecting the total amount of protein inside the cell (Fig. 1C). The amount of protein varied considerably among the dynein and regulatory proteins. Next, we quantified the average intensity at the nuclear/spindle region (nuclear intensity or NI), and the value at the brightest time point was plotted (Fig. 1D). The variation in the CI was roughly preserved for the NI with one exception, i.e., the NI of NUD-2 protein increased to a level comparable to that of dynein and dynactin, while the CI of NUD-2 was the lowest among the proteins investigated here. This exception was evident when we calculated the N/C ratio, the value obtained by dividing the NI by the CI (Fig. 1E). As expected, NUD-2 showed a higher degree of enrichment compared to the other proteins. This suggested the possibility that NUD-2 accumulated with a specific mechanism to help achieve a concentration comparable to that of dynein. Another interesting feature of the N/C ratio, except NUD-2, was that it was almost constant, while the total amount of the proteins (CI) varied. A simple explanation might be that these proteins shared a common import/export mechanism, and thus that the equilibrium ratio was constant.

### Variations in the target sites of accumulations among the proteins

To further investigate the nature of accumulation of dynein and its regulatory proteins, we focused on the spatial distribution of the accumulated proteins. We assumed three candidate sites for accumulation in the spindle regions, namely kinetochores, spindle microtubules, and bulk spindle regions (**Figure S2A**). The former two sites are well-known associated regions of dynein and several regulatory proteins (Heald and Khodjakov, 2015). Owing to the abundance of microtubules at the spindle region, it could not be easily ascertained whether a protein was bound to microtubules or whether accumulation occurred at other sites in the spindle. To eliminate the effects of contribution of microtubules, we used nocodazole treatment and observed the embryos with microtubules depolymerized. In the nocodazole-treated embryos, LIS-1, NUD-2, and LIN-5 continue to demonstrate evident accumulation around the time of CeNEBD, an event which has been reported to correspond to the accumulation at the nascent spindle region (**Figures 2A-C and Movie S2**). LIS-1 and NUD-2 later accumulated around the chromosomes (**Figures 2A and B**), while LIN-5 was excluded from the chromosome region (**Figure 2C, arrowheads**). As the *C. elegans* chromosome is holocentric, exhibiting possession of multiple kinetochores along the entire length of the chromosome, the localization of LIS-1 around the chromosome has been assumed to be associated with kinetochores, a finding which is consistent with that reported in a previous study (Cockell *et al.*, 2004; Simões *et al.*, 2018).

**Figure 2.**
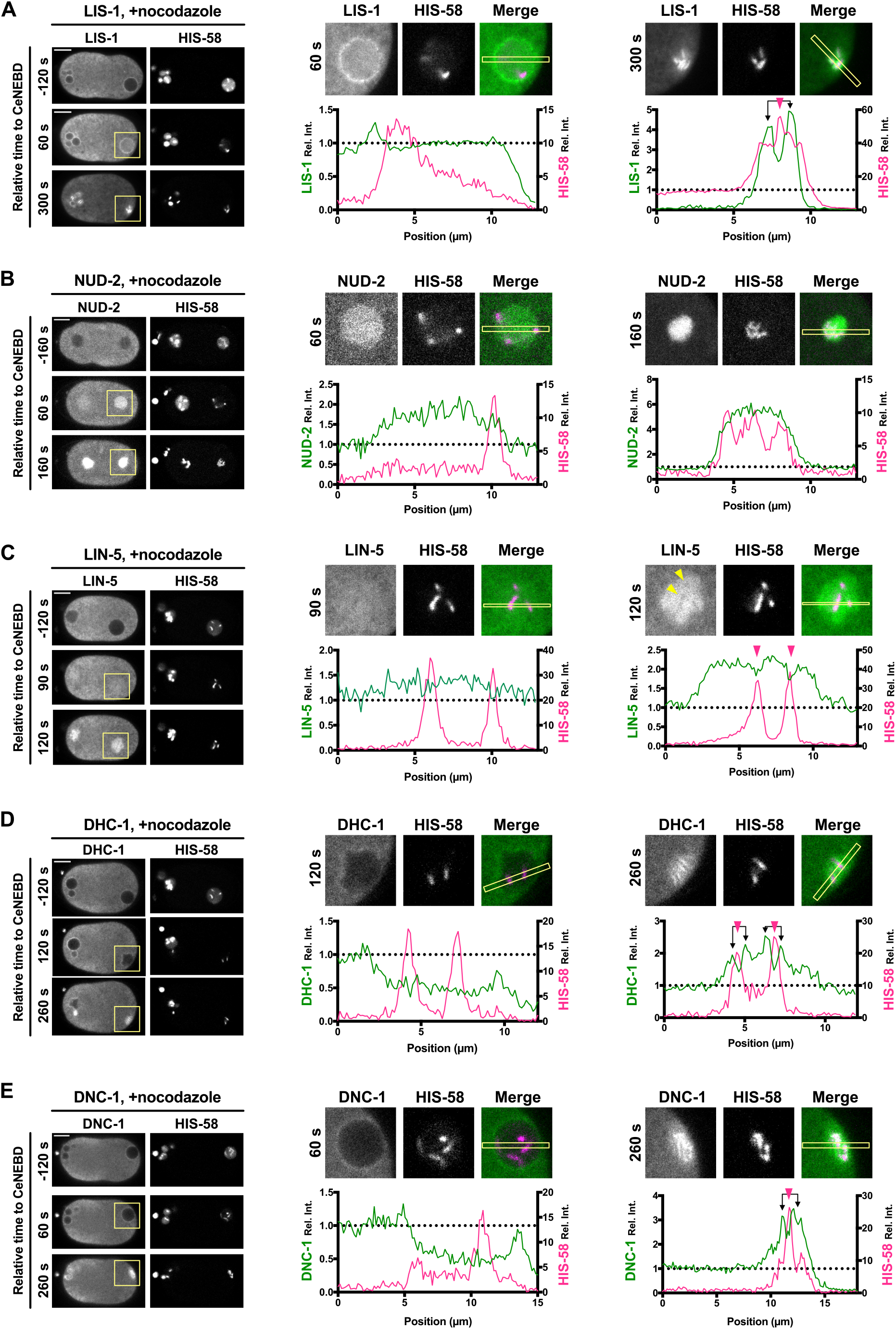
Variations in accumulation sites of dynein and its regulatory proteins. Spatial distribution of LIS-1 (A), NUD-2 (B), LIN-5 (C), dynein (D), and dynactin (E), in the presence of 10 μg/mL of nocodazole are presented by depicting the single-plane time-lapse images (left), the magnified images (center), and the intensity profiles (right). Loss of microtubules via nocodazole treatment leads to the defects in pronuclear migration and meeting because these processes are mediated by microtubule-based motors. It results in the delay in CeNEBD of oocyte pronuclei due to the lack of triggering signal arising from centrosome-associated molecules. The left side of the image corresponds to the anterior. The magnified images have been cropped from the yellow rectangles in the images depicting the corresponding time. Intensity profiles have been calculated in the rectangles indicated in the magnified images. The magenta arrowheads denote the peak of histone signals, and the black arrow lines indicate the signal peak of dynein or the regulatory proteins near the histone peaks. The scale bars indicate 10 μm.

In contrast, dynein and dynactin did not show apparent accumulation in the bulk spindle region, indicating that spindle microtubule accumulation at the spindle shown in Figure 1A was mediated by spindle microtubules. Under the nocodazole-treated condition, late accumulation was observed at the chromosomes (**Figures 2D and 2E, and Movie S2**). Such findings on chromosomal accumulations were consistent with those reported in previous studies (Gassmann *et al.*, 2008; Bader and Vaughan, 2010). This finding indicated that both proteins accumulated in the spindle region through the establishment of interaction with kinetochores and spindle microtubules. These spatial patterns of accumulation were also confirmed in the *tbb-2* (*RNAi*) embryos, where tubulin expression was impaired (**Figure S2B**). Our results indicated that the proteins showed a spatial variation in their accumulation, and a few proteins could accumulate independently of spindle microtubules. In terms of dynein regulation, the results suggest that accumulation in the nascent spindle region before dynein recruitment may contribute to the efficient formation of the required complex.

### Variations in the timing of accumulations

In our previous analyses, we showed that tubulin accumulated in the nascent spindle region with the occurrence of CeNEBD and suggested that other proteins could also enter the region upon CeNEBD (Hayashi *et al.*, 2012). In the present study, however, we observed that NUD-2 entered and accumulated in the nuclear region before NEBD (**Figure 3A**). Interestingly, NUD-2 is unique among dynein regulatory proteins because of its unique timing of accumulation and high N/C ratio (**Figure 1B**). The accumulation mechanism of NUD-2 has been discussed in detail in later sections. Inspired by the early accumulation events of NUD-2 and the distinct localization pattern in the spindle region (**Figure 2**), we investigated the accumulation timing of dynein and the regulatory proteins comprehensively. We analyzed the time series of the N/C ratio of dynein and the regulatory proteins in the 1-cell stage *C. elegans* embryos. The time series of the N/C ratio indicated that these proteins did not accumulate in the spindle region simultaneously (**Figure 3A**). NUD-2 accumulated earliest among the proteins, followed by LIS-1 and LIN-5. After the accumulation of tubulin, dynein and dynactin accumulated at similar times. This finding was consistent with that of the spatial analysis described above, which suggested that dynein and dynactin accumulated mainly through the establishment of interaction with microtubules. There were no apparent differences in the accumulation rates among the proteins.

**Figure 3.**
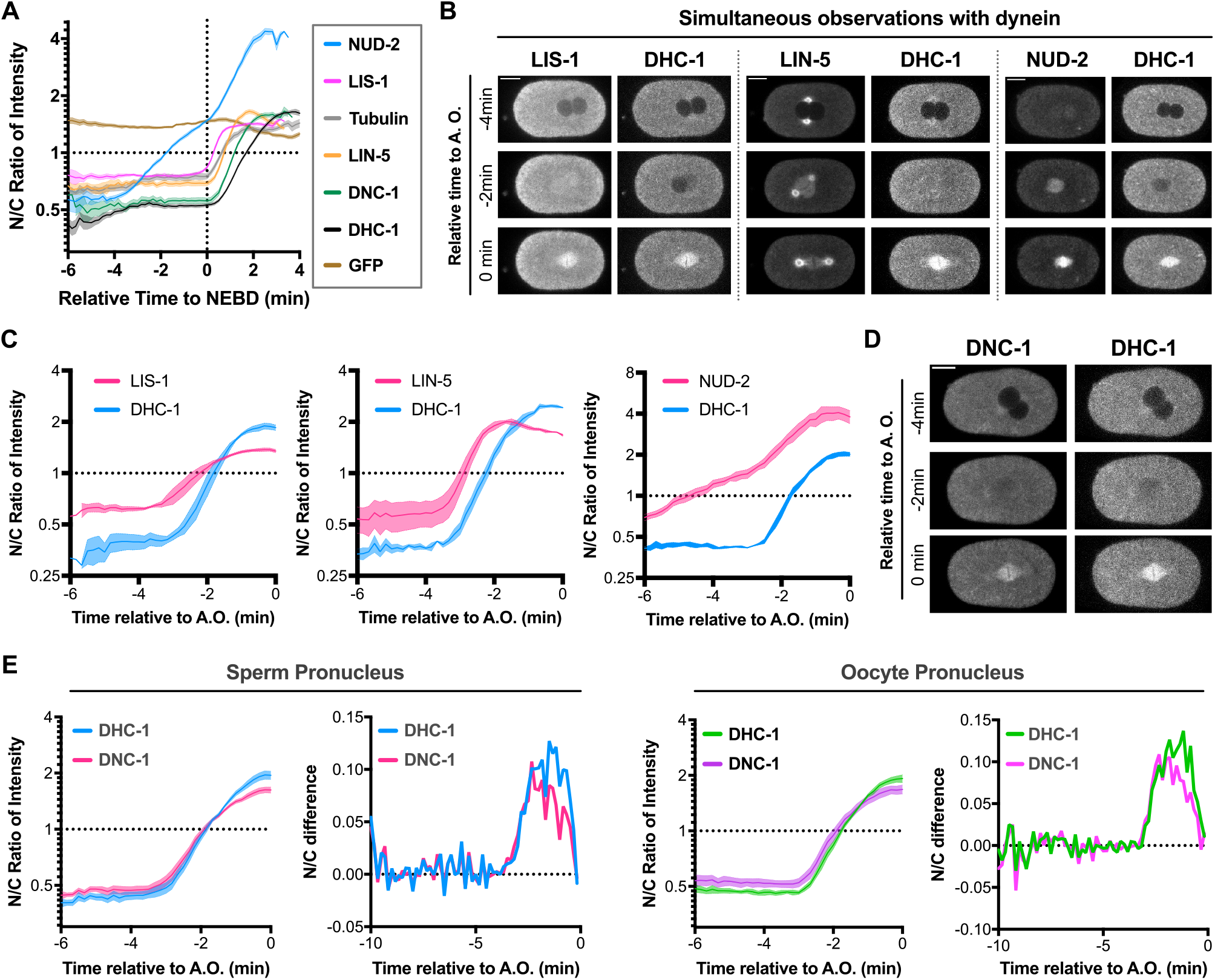
Temporal variations in the accumulation. (A) The plots show the time series of the intensity ratio in the spindle regions to the cytoplasm (N/C ratio). The initiation time of CeNEBD, the time origin, has been determined using the intensity decay of free histone (HIS-58) in the nucleus. The numbers of nuclei analyzed were 16 from 8 embryos (NUD-2), 19 from 11 embryos (LIS-1), 16 from 12 embryos (tubulin), 15 from 10 embryos (LIN-5), 18 from 11 embryos (dynactin), and 26 from 14 embryos (dynein). The mean and SE values are shown. (B and C) Typical single-plane time-lapse images (B) and time series of the N/C ratio (C) were obtained via the simultaneous observations of dynein with LIS-1, LIN-5, and NUD-2. (C) The right side of the image corresponds to the anterior. The scale bars indicate 10 μm. Owing to the lack of information on histone signal, the time origin was set to anaphase onset, which was determined via the segregation of chromosome- localized signals of dynein. (C) The N/C ratio used to perform calculations based on sperm pronuclei is shown. The number of nuclei analyzed was 4 from 4 embryos (LIS-1), 3 from 3 embryos (LIN-5), and 4 from 4 embryos (NUD-2). (D and E) Typical single-plane time-lapse images (D) and time series of the N/C ratio (E) obtained via the simultaneous observation of dynein and dynactin. (E) The plots show the time series of the N/C ratio and its time difference. The numbers of nuclei analyzed were 8 from 8 embryos (sperm pronuclei) and 7 from 8 embryos (oocyte pronuclei).

Although our observations suggested a sequential accumulation pattern of dynein and the regulatory proteins, it has been demonstrated based on experiments conducted using different strains expressing each protein fused to the fluorescent protein. To further investigate whether the timing of accumulation of the two proteins was the same, we conducted simultaneous observations of dynein and the regulatory proteins by constructing the strains simultaneously expressing the two fluorescently labeled proteins (**Figure 3B and Table S1**). We found that NUD-2, LIS-1, and LIN-5 accumulated earlier than dynein, as suggested by previous observations, whereas dynactin showed approximately the same timing of accumulation (**Figures 3B and 3C**). In the case of LIN-5, NUD-2, and LIS-1, the simultaneous observations revealed marked differences in accumulation patterns, suggesting that these proteins were not associated with dynein upon entry into the spindle region. The simultaneous accumulation of dynein and dynactin suggested that a considerable fraction of the proteins demonstrated association with each other in the cytoplasm and presented with transportation as complexes. Interestingly, although dynein and dynactin gradually accumulated almost simultaneously, they displayed a different pattern around the time of saturation; when the accumulation speed of dynactin decreased, dynein continued to exhibit a maximum accumulation speed (**Figure 3C**). These results suggest that at least a certain proportion of the accumulated dynein did not form a complex with dynactin.

### Molecular weight is not the determinant of accumulation order

We observed that the timing of accumulation differed between dynein and its regulatory proteins. The proteins were expected to enter the spindle region mainly through diffusion because NPCs, which act as a diffusion barrier and as a mediator for active nucleocytoplasmic transport, underwent disassembly by that time (Tzur and Gruenbaum, 2013). Thus, we hypothesized that the difference in diffusion rate depending on molecular weight accounted for the temporal difference. This hypothesis was supported by the fact that the accumulation order of dynein-regulatory proteins coincided with the order of molecular weights; NUD2 (~69 kDa) accumulated first, followed by LIS-1 (92 kDa) and LIN-5 (187 kDa), with final accumulation of dynactin and dynein (>1.2 M) (**Figure 3A**). To examine the effect of molecular weight on the accumulation, we observed the temporal dynamics of polymers with defined molecular sizes using an injection method (**Figure S3A**) (Galy *et al.*, 2003; Updike *et al.*, 2011). We incorporated polymers of different sizes into *C. elegans* embryos through oogenesis and compared the temporal dynamics.

Although previous studies have reported the presence of injected dextran in interphase embryos (Galy *et al.*, 2003; Updike *et al.*, 2011), it was unclear whether they accumulated at the spindle region during mitosis. Thus, we decided to observe the accumulation events of polyethylene glycol (PEG) as well as dextran. By observing dextran (40 kDa) and PEG (40 kDa) accumulations, we found that dextran showed CeNEBD-dependent accumulation in the spindle region (**Figure 4A**), while PEG was excluded from the nucleus throughout the cell cycle (**Figure 4B**). Notably, PEG with a smaller molecular weight (5 kDa), which was expected to be below the diffusion limit of NPCs, was also excluded from the nucleus (**Figure S3B**), suggesting that the event of accumulation of a polymer at the spindle region was dependent on physicochemical properties, such as the existence of branching in the polymer structure.

**Figure 4.**
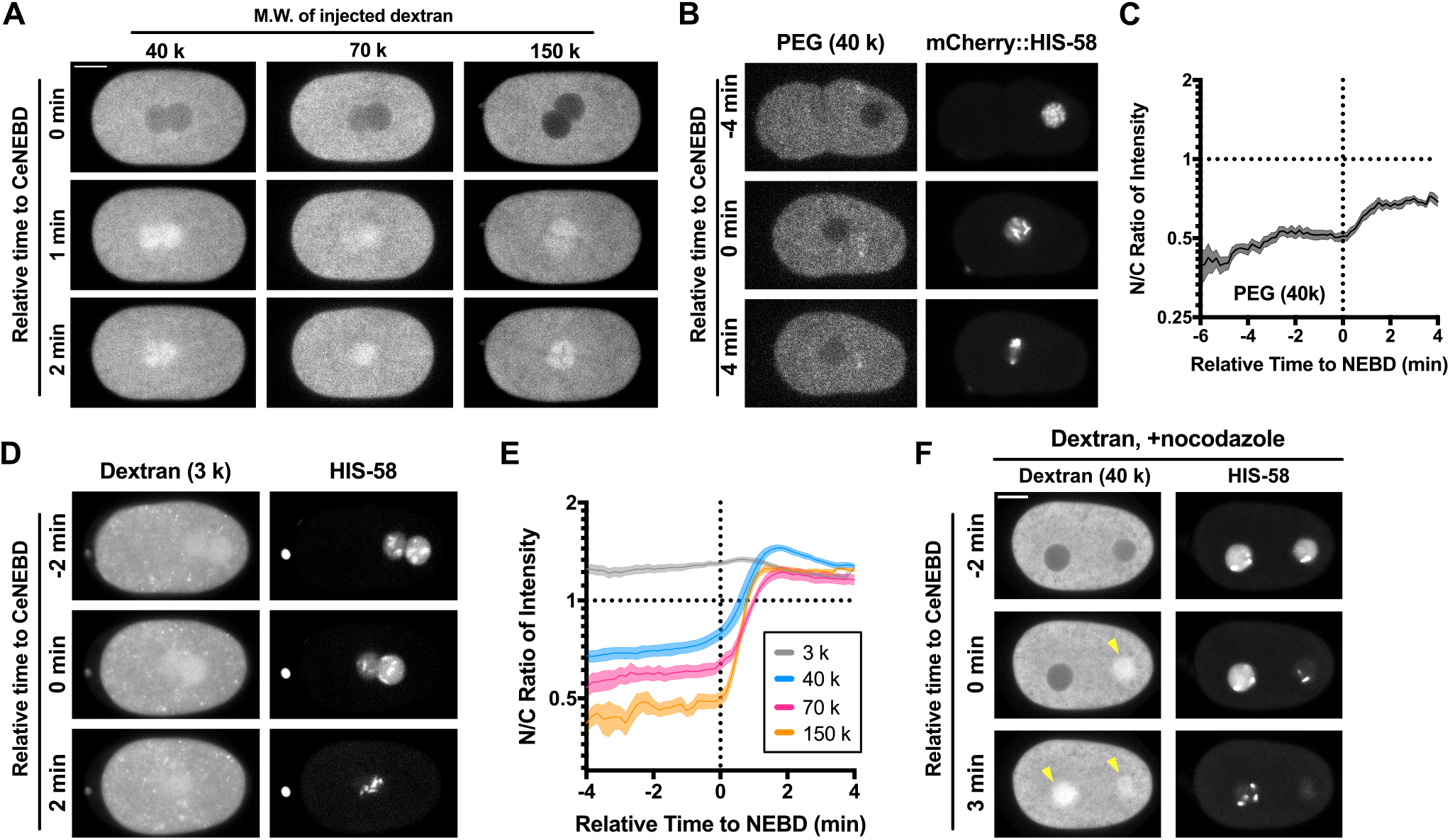
Accumulation dynamics of polymers incorporated into the embryos. (A) Typical single-plane time-lapse images showing the temporal dynamics of dextrans incorporated through the gonad injection. The molecular weight of the injected dextran is indicated above. The left side of the image corresponds to the anterior. The scale bars indicate 10 μm. (B) Typical single-plane time-lapse images showing the temporal dynamics of mPEG (40 k). The left side of the image corresponds to the anterior. The scale bar indicates 10 μm. (C) Time series of the N/C ratio of mPEG (40 k). The number of analyzed pronuclei was 14 from 10 embryos. Mean and SEM are shown. (D) Typical single-plane time-lapse images showing the temporal dynamics of dextran (3 k). The left side of the image corresponds to the anterior. (E) Time series of the N/C ratio of the injected dextrans (3k, 40 k, 70 k, and 150 k). The numbers of pronuclei analyzed were 9 from 6 embryos (3 k), 7 from 4 embryos (40 k), 8 from 4 embryos (70 k), and 9 from 6 embryos (150 k). Mean and SEM are shown (F) Typical single-plane time-lapse images showing the accumulation of dextran (40 k) in the presence of 10 μg/mL nocodazole. The scale bar indicates 10 μm. The left side of the image corresponds to the anterior.

We then compared the accumulation dynamics of dextrans with molecular weights of 3, 40, 70, and 150 k. Dextran (3 kDa) presented with continuous accumulation in the nuclear region throughout the cell cycle (**Figure 4D**), probably because the molecular weight was below the diffusion threshold of the nuclear pore complex (**Figure 4D**). Other dextrans exhibited CeNEBD- dependent accumulation at the nascent spindle region (**Figure 4A and Movie S3**). As depicted in the time series based on the N/C ratios, dextrans accumulated only after CeNEBD (**Figure 4E**). The time series also did not demonstrate any marked difference in the timing of dextran accumulation (**Figure 4E**); at least the difference could not be considered to explain the difference between LIS-1 (92 kDa) and LIN-5 (187 kDa) by molecular weight (**Figure 3A**). This result indicated that molecular weight was not a determinant factor for the accumulation order.

Although molecular weight was not deemed the determinant, it was notable that exogenous polymers showed an accumulation pattern similar to that shown by dynein and the regulatory proteins. Additionally, it was observed that the proteins, dynein and dynactin, mainly accumulated through the establishment of interaction with microtubules (**Figures 2D and 2E**). Thus, we examined whether the accumulation of dextran depended on the interaction with microtubules. The observation of dextran in the nocodazole-treated embryos showed that it continued to accumulate at the nascent spindle region (**Figure 4F**), indicating that the accumulation of dextran was not dependent on microtubules such as LIS-1, NUD-2, and LIN-5 (**Figures 2A-C**). Furthermore, similar to LIN-5, dextrans showed a uniform distribution in the nascent spindle region (**Figure S3D**). Although the accumulation dynamics of dextrans shared several characteristics with dynein and the regulatory proteins, dextran did not present with accumulation before CeNEBD, as that observed for NUD-2, suggesting an additional requirement for such an accumulation pattern.

### Pre-NEBD accumulation of NUD-2 is independent of CeNEBD

Among the proteins observed, NUD-2 showed a distinct accumulation pattern compared with the other proteins; accumulation started approximately 4 min before CeNEBD and the highest maximum N/C ratio of approximately 4.5-fold was noted (**Figures 1B and 3A**). This pronounced accumulation before CeNEBD was observed only for NUD-2, and to our knowledge, this was a unique phenomenon. We termed this phenomenon “pre-NEBD accumulation” and investigated it comprehensively.

We observed that the initiation time of pre-NEBD accumulation was around the time of the pronuclear meeting. If the pre-NEBD accumulation is dependent on pronuclear meetings, it should occur only at the 1-cell stage because the pronuclear meeting is specific to the 1-cell stage. However, this was not the case. We found that pre-NEBD accumulation also occurred in the later stage embryos (2–16-cell stage; **Figure 5A**). Interestingly, as development proceeded, the degree of accumulation through pre-NEBD accumulation increased, while the final N/C ratio after post-NEBD accumulation did not vary among the cell stages (**Figures 5B-D**). In contrast to the early embryos, in oocyte, NUD-2 did not accumulate to the nuclear region prior to the NEBD of the oocyte meiosis. The post-NEBD accumulation was observed for the oocyte meiosis (**Figures 5E and 5F**). Moreover, we found that NUD-2 localized at the nuclear membranes in all oocytes except the most proximal (−1) one (**Figure 5E**), in contrast to the early embryos. These results suggest that pre-NEBD accumulation is specific to mitotic division, whereas post-NEBD accumulation is universal to mitosis and meiosis.

**Figure 5.**
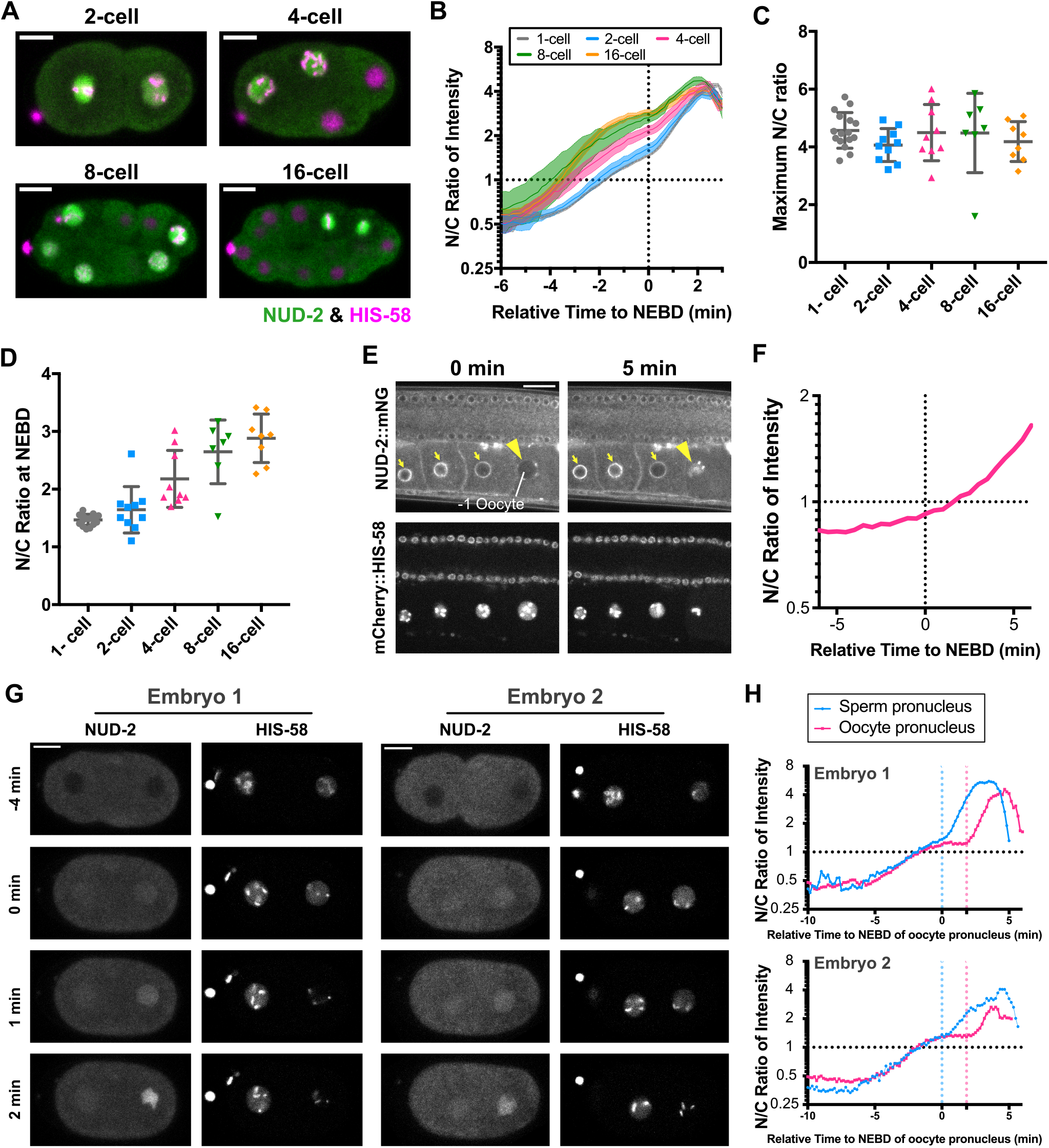
Accumulation patterns of endogenous NUD-2 in various contexts. (A) Single plane time-lapse images showing the signal of NUD-2::mNG (green) and mCherry::HIS-58 (magenta) in 2–16-cell stage embryos. The right side in the images corresponds to the anterior. The scale bars indicate 10 μm. (B) Time series of the N/C ratio of NUD-2 in 2-cell (blue), 4-cell (red), 8-cell (green), and 16-cell (orange) embryos. For comparison, the time series of NUD-2 in 1-cell stage embryos is shown by using the gray line, which indicates the same data as indicated in Figure 2. (C) Maximum N/C ratio in 1–16-cell stage embryos. (D) The N/C ratio measured at the time of CeNEBD. (E) Typical single plane time-lapse images depicting NUD-2 in the germline of an adult worm. The yellow arrowheads denote the nucleus of the - 1 oocyte, and the yellow arrows indicate the NUD-2 localizations at nuclear membranes. The times relative to CeNEBD of the −1 oocyte are indicated above. The scale bar indicates 20 μm. (F) Time series of the N/C ratio of NUD-2 in the −1 oocyte shown in (E). (G) Typical single-plane time-lapse images showing the temporal dynamics of NUD-2 in the presence of 10 μg/mL nocodazole. The scale bars indicate 10 μm. (H) Time series of N/C ratio of NUD-2 in the nocodazole-treated embryos. The N/C ratios in sperm and oocyte pronuclei are depicted by using the blue lines and the magenta lines. The vertical dashed lines indicate the initiation of CeNEBD of pronuclei.

We then investigated the relationship between pre-NEBD accumulation of NUD-2 and CeNEBD. We focused on a key aspect: was pre-NEBD accumulation coupled with CeNEBD? If pre- NEBD accumulation depends on CeNEBD, the timing of pre-NEBD accumulation between sperm and oocyte pronuclei will differ in the presence of nocodazole. Nocodazole treatment impairs pronuclear meeting, which in turn delays NEBD of the oocyte pronucleus due to the lack of signals from centrosomes attached to the sperm pronucleus (Hachet *et al.*, 2007; Portier *et al.*, 2007; Toya *et al.*, 2011). When NEBD of oocyte pronucleus was delayed, there was no delay in the initiation time of pre-NEBD accumulation and it occurred at the same time as that of sperm pronucleus (**Figures 5G and 5H**). After reaching a value of approximately 1.3, the N/C ratio of the oocyte pronuclei showed the achievement of a steady state for several minutes, while the N/C ratio of the sperm pronucleus showed a transition to post-NEBD accumulation. These results suggest that pre-NEBD accumulation is a distinct process from the post-NEBD accumulation and is regulated by factors independent of CeNEBD.

### NUD-2 exhibits a distinct accumulation pathway from tubulin

Ran, a small GTPase protein, plays a central role in nuclear transport. A recent study revealed that Ran contributed to the accumulation of a tubulin chaperone in the nuclear region before NEBD in *Drosophila melanogaster* (Métivier *et al.*, 2021). We have previously shown that Ran-1 is necessary for the post-NEBD accumulation of tubulin in *C. elegans* embryos (Hayashi *et al.*, 2012). We sought to ascertain whether Ran was involved in the pre-NEBD accumulation of NUD-2 by conducting knockdown experiments for *C. elegans* Ran, *ran-1*. In RAN-1-depleted embryos, we confirmed a reduction in cell size, nuclear size, and observed defects in mitosis (**Figure 6A**), as those previously described (Gönczy *et al.*, 2000; Askjaer *et al.*, 2002; Bamba *et al.*, 2002). The defect in cytokinesis maintained the embryos in the 1-cell stage, although the nuclei underwent multiple divisions. Even under such conditions, we observed the cyclic accumulation of NUD-2 at the sites of histone signals (**Figures 6A and 6B, and Movie S4**). Such accumulation was not observed for tubulin (**Figures 6C and 6D**), consistent with the finding reported in a previous study, which revealed the contribution of RAN-1 to tubulin accumulation (Hayashi *et al.*, 2012). These results suggest that NUD-2 exhibits a different accumulation pathway from tubulin, whose post-NEBD accumulation is dependent on RAN-1 (Hayashi *et al.*, 2012).

**Figure 6.**
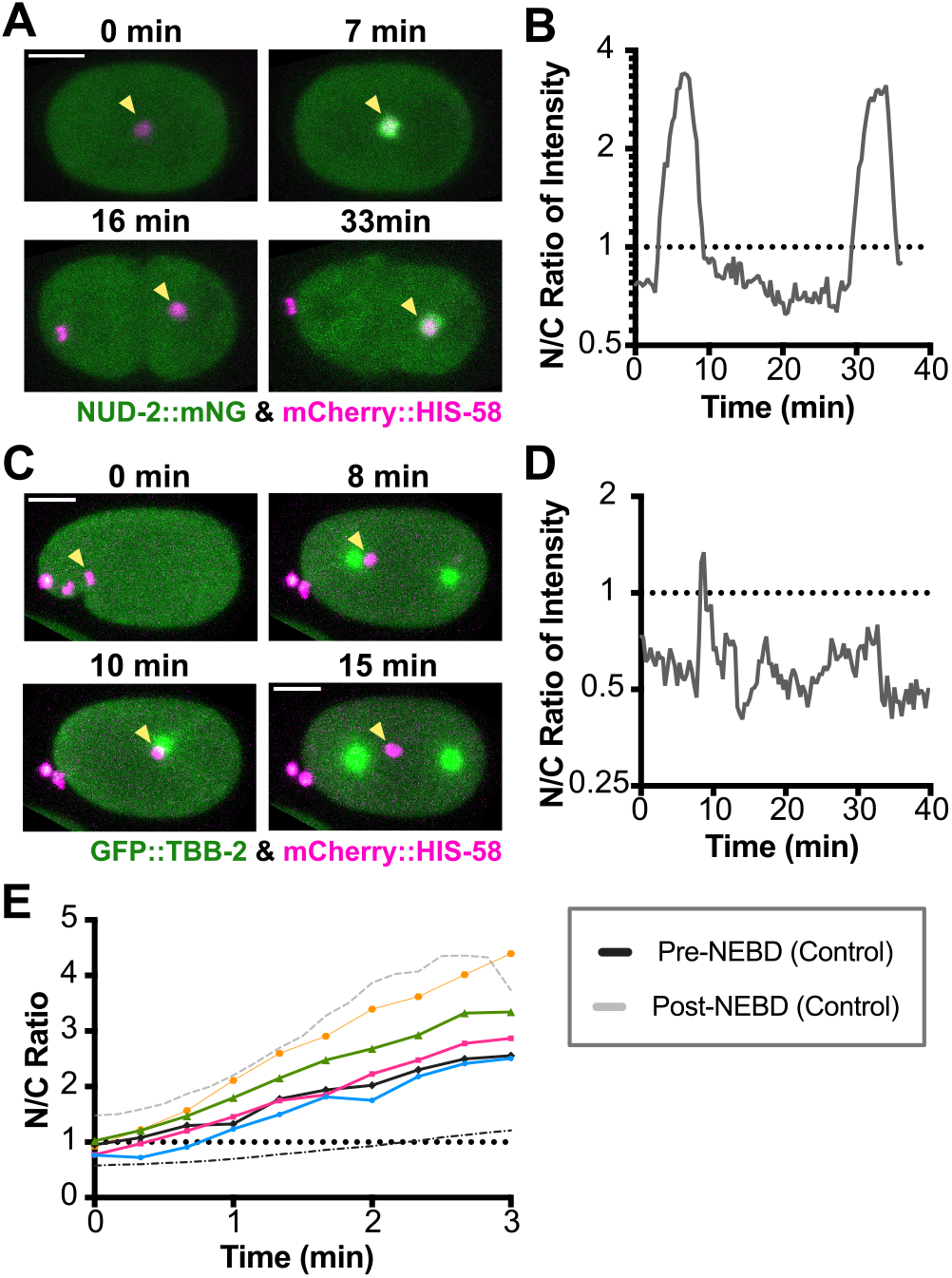
Accumulation of NUD-2 exhibits a distinct molecular dependency from tubulin. (A) Maximum projection images showing the temporal dynamics of NUD-2 in the RAN-1-depleted embryo. Under *ran-1* (RNAi) conditions, the sizes of embryo and nucleus reduced, and the defect in cytokinesis was observed. Although it was difficult to detect the precise timing of CeNEBD, the cyclic increase in NUD-2 signals was confirmed. The left side of the image corresponds to the anterior. The scale bar indicates 10 μm. (B) Time series of N/C ratio of NUD-2 in the RAN-1-depleted embryos shown in (A). The intensity of NUD-2 in the region indicated by using the yellow arrowheads was measured. The origin of time was set to the initial time of the observation. Each peak seemed to demonstrate a rapid single-phase increase from the N/C ratio below 1. (C) Maximum projection images showing the temporal dynamics of TBB-2 (tubulin) in the RAN-1- depleted embryo. The left side of the image corresponds to the anterior. (D) Time series of N/C ratio of TBB- 2 in the RAN-1-depleted embryos shown in (C). The intensity of NUD-2 in the region indicated by using the yellow arrowheads was measured. The origin of time was set to the initial time of the observation. (E) Comparison of temporal dynamics of the N/C ratio between the unperturbed condition and ran-1 (RNAi) conditions. The time series data of N/C ratio during the rapid increase phase in *ran-1* (RNAi) embryos are shown by using the colored lines, while the black line and the gray line show the time series data of pre- and post-NEBD accumulations, respectively. The origin of time was set as the initial time of each accumulation, not the timing of CeNEBD because it was difficult to detect CeNEBD in the RAN-1-depleted embryos.

To investigate the details of NUD-2 accumulation in RAN-1-depleted embryos, we analyzed the time series of the N/C ratio. In *ran-1* (RNAi) embryos, we could not determine the timing of NEBD from the localization pattern of histones, and thus it was difficult to differentiate between pre-NEBD and post-NEBD accumulation events of NUD-2. As shown in **Figure 6E**, NUD-2 signals increased at an approximately constant rate. This increasing pattern was somewhat different from the unperturbed condition where we observed slower accumulation followed by a short constant phase before NEBD and faster accumulation after NEBD (**Figure 3A**). We considered that either pre- or post-NEBD accumulation was impaired by *ran-1* (RNAi). By comparing the rate of accumulation, we found that the accumulation rate under the *ran-1* (RNAi) condition was more similar to that under the unperturbed condition (**Figure 6E**). Furthermore, the maximum N/C ratio of NUD-2 in the absence of RAN-1 was estimated to be 3.8 ± 1.6 (based on 7 increase events in 5 embryos), comparable to that of post-NEBD accumulations under the unperturbed condition (4.7 ± 0.6). These results suggest that Ran is necessary for pre-NEBD accumulation, but is not vital in the post-NEBD accumulation of NUD-2. This is in contrast to tubulin, where post-NEBD accumulation is impaired by *ran-1* (RNAi) (Hayashi *et al.*, 2012). These results suggest that the mechanism of post-NEBD accumulation is different between NUD-2 and tubulin.

### The spindle region requires NUD-2 to retain the accumulated proteins

We noted a distinct pre-NEBD accumulation timing for NUD-2 compared with other proteins. The earliest accumulation and the highest N/C ratio of NUD-2 suggested a role for NUD-2 in the accumulation of other proteins in the spindle region. We observed the accumulation pattern of dynein and the regulatory proteins LIS-1, dynactin, and LIN-5 in NUD-2-depleted embryos. Even in the absence of accumulated NUD-2, post-NEBD accumulation continued to occur (**Figure 7A**). However, their localization level at the spindle was lower than that under the unperturbed condition for all proteins except LIN-5 (**Figures 7A and 7B**). A previous study using the deletion mutant of NUD-2 has already reported the reduction in LIS-1, dynactin, and dynein at the kinetochore (Simões *et al.*, 2018), and our observations showed that the reduction also occurred along the entire spindle. To further investigate the effect of NUD-2 depletion, we analyzed the time series of the N/C ratio. The derived time series indicated that LIS-1, dynactin, and dynein showed decays in the N/C ratio after the initial phase of increase (**Figure 7C**). This finding suggested that the ability of the bulk spindle region to retain the accumulated proteins was impaired in NUD-2-depleted embryos. In addition to the effect on the temporal dynamics of dynein-regulatory proteins, a previous study reported chromosome abnormalities in the deletion mutant of *nud-2* (Simões *et al.*, 2018). We examined chromosome dynamics in NUD-2-depleted embryos and confirmed abnormalities in chromosome dynamics (**Figure S4**). Approximately half of the embryos showed lagging chromosomes (13/26 embryos), and more than 80% of the embryos presented with additional histone signals (20/26 embryos). These abnormalities were consistent with those reported in the previous study (Simões *et al.*, 2018) and were probably caused by meiosis/mitosis defects (**Figure S5**). Since dynein has been reported to be involved in chromosome alignment and segregation (Bader and Vaughan, 2010; Raaijmakers and Medema, 2014), it was difficult to determine whether the chromosomal abnormalities were attributed to the direct or indirect consequences of NUD-2 depletion. A decrease in the amount of dynein or activation complex in the spindle region may also be related to abnormalities as an indirect effect.

**Figure 7.**
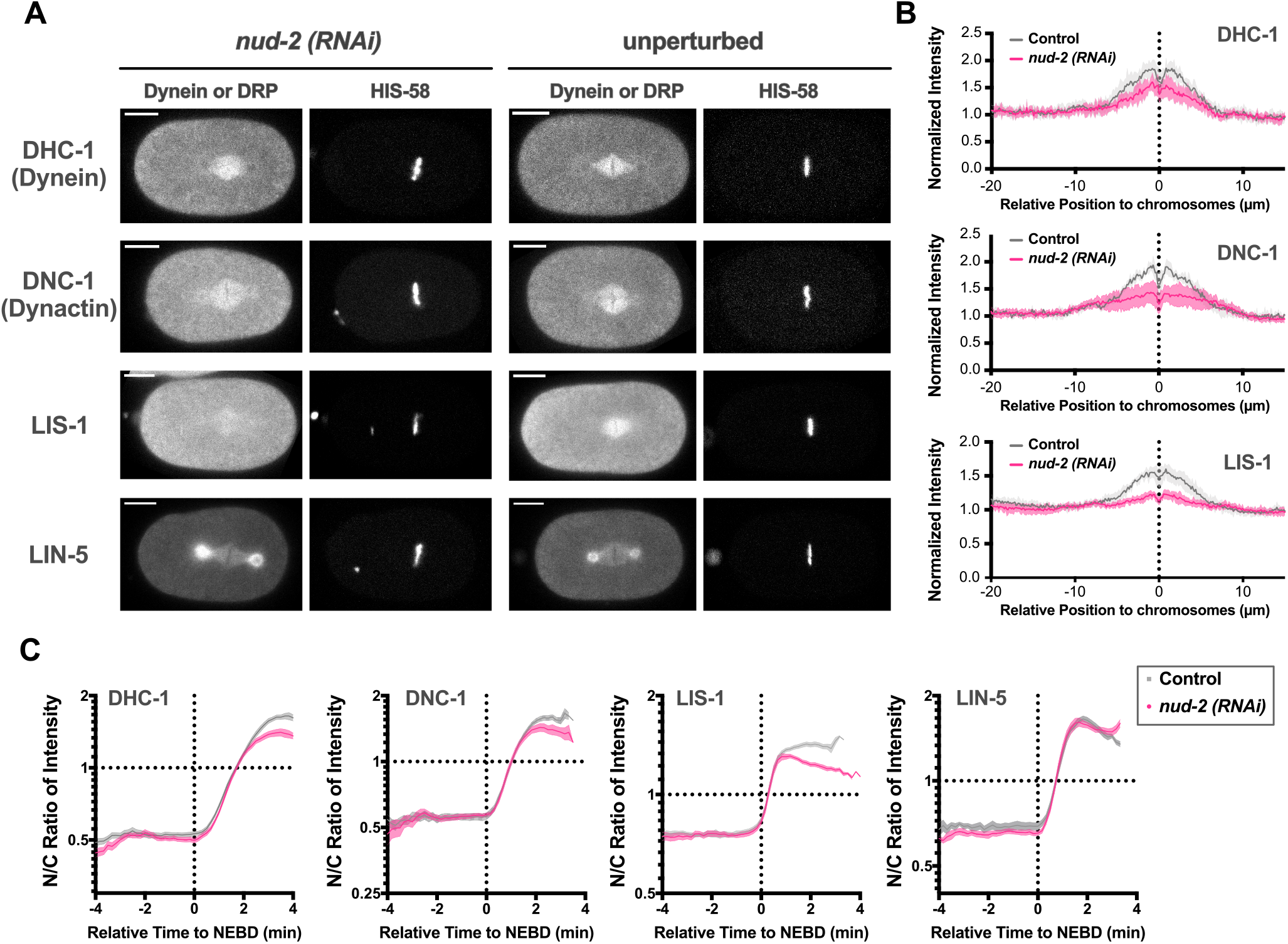
NUD-2 depletion reduces the retained amount of dynein and its regulatory proteins. (A) Typical single-plane images of dynein, dynactin, LIS-1, and LIN-5 showing the comparison between the nud-2 (RNAi) embryos and the unperturbed embryos. The images of embryos at the anaphase onset were acquired. The left side of the image corresponds to the anterior. The scale bars indicate 10 μm. (B) The averaged intensity profiles of dynein (top), dynactin (middle), and LIS-1 (bottom) were derived from both the unperturbed (control) and the nud-2 (RNAi) embryos. The profiles measured before the anaphase onset are shown. The magenta lines indicate the profiles derived from the nud-2 (RNAi) embryos, whereas the gray lines indicate the profiles derived from the control embryos. For all analyzed proteins, a reduction in intensity was observed around the chromosomes. (C) Temporal dynamics of N/C ratio of dynein and its regulatory proteins under the unperturbed or nud-2 (RNAi) conditions. The gray dots indicate the time series under the unperturbed conditions, whereas the magenta dots indicate the time series in nud-2 (RNAi) embryos. Before the initiation of a rapid accumulation phase occurred around CeNEBD, there was no evident difference between the controls and the nud-2 (RNAi) conditions. All analyzed proteins except LIN-5 showed a decay of N/C ratio after the rapid accumulation. The numbers of nuclei analyzed were 22 from 12 embryos (dynein), 13 from 8 embryos (dynactin), 13 from 7 embryos (LIS-1), and 15 from 9 embryos (LIN-5).

### The C-terminal helix region of NUD-2 is responsible for pre-NEBD

NDEL1/NDE1, the human ortholog of NUD-2, possesses two distinct structural regions. The N-terminal region forms an approximately 20 nm-long coiled-coil structure (Derewenda *et al.*, 2007), whereas the C-terminal region is predicted to be intrinsically disordered. Within the C-terminal region, NDEL1/NDE1 possesses a putative helix region flanked by two intrinsically disordered sequences. The N-terminal region includes one of the two dynein-binding sites and the LIS-1 binding site (Derewenda *et al.*, 2007), aiding binding between LIS-1 and dynein (Zyłkiewicz *et al.*, 2011; Wang *et al.*, 2013). The C-terminal region contains a second dynein-binding site and many phosphorylation sites (Niethammer *et al.*, 2000; Yan *et al.*, 2003; Mori *et al.*, 2007; Bradshaw *et al.*, 2008). Based on these structural and functional findings, we investigated the region of NUD-2 necessary for the characteristic pre-NEBD accumulation. For this purpose, we adopted a protein-injection approach. We injected the recombinant NUD-2 fragments into the *C. elegans* gonad and observed the temporal dynamics of the fragments incorporated into the embryos through oogenesis (**Movie S5**). Before observing NUD-2 fragments, we validated our injection method by examining the temporal dynamics of mCherry, which was consistent with that of the transgenic GTP (**Figure S5**).

We first observed accumulation of injected full-length NUD-2. Compared to the endogenous protein (Figures 1A and 3A), although the injected full-length NUD-2 accumulated at the spindle region, it showed a lower maximum N/C ratio and slower accumulation rate, which resulted in an N/C ratio below 1 at CeNEBD (Figure 8C). However, we found that the initiation time of accumulation was earlier than that of CeNEBD, and there was a change in the accumulation rate around CeNEBD (Figure 8C). Based on these results, we concluded that the injected protein showed both pre- and post-NEBD accumulation, although the accumulation rate was less than the endogenous one. The reduction in accumulation might be attributed to the saturation of accumulation caused by the presence of markedly more NUD-2 level than the control. We then observed the N- terminal fragment composed of the predicted coiled-coil domain (NUD-2_CC_, 2-165 aa). In contrast to the full-length construct, NUD-2_CC_ was excluded from the pronuclei until CeNEBD, without pre- NEBD accumulation (Figures 8B and 8C). We then observed the C-terminal fragment (NUD-2_IDR_, 166-293 aa) and found that NUD-2_IDR_ showed both pre- and post-NEBD accumulations. To further investigate which region was responsible for pre-NEBD accumulation, we next observed NUD-2_CC-IDR1_ (2-238 aa), including the coiled-coil and the following intrinsically disordered regions. NUD-2_CC-IDR1_ was excluded from the interphase nucleus and only accumulated after CeNEBD, which exhibited the same temporal pattern as NUD-2_CC_. Finally, we observed NUD-2_CC-IDR1-H_ (2-276 aa), which contained the region comprising the N-terminal coiled-coil and the C-terminal helix. Interestingly, NUD-2_CC-IDR1-H_ showed both pre- and post-NEBD accumulations. These results suggest that the C-terminal helix region is involved in the pre-NEBD accumulation of NUD-2.

**Figure 8.**
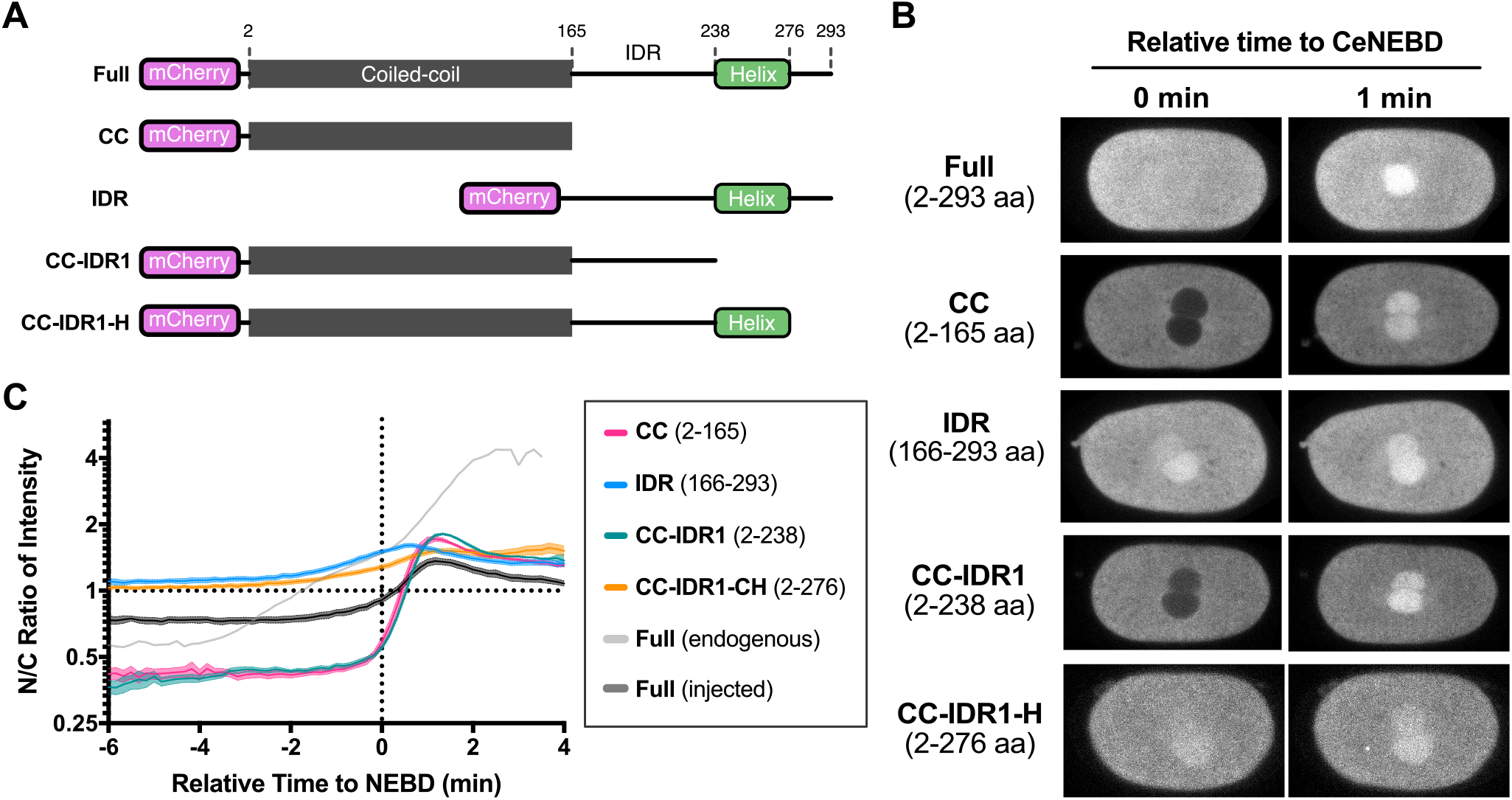
The C-terminal helix of NUD-2 is necessary for pre-NEBD accumulation. (A) Schematic representation of the recombinant NUD-2 proteins used for the injection experiments. The fragments were purified from *E. coli* cells. The predicted coiled-coil region is depicted as gray boxes. IDR indicates an intrinsically disordered region. (B) Typical single-plane time-lapse images showing the temporal dynamics of the NUD-2 fragments. The left sides of the images correspond to the anterior. The scale bars indicate 10 μm. (C) Time series of the N/C ratio of the recombinant NUD-2 proteins. The numbers of nuclei analyzed were 10 from 6 embryos (Full), 9 from 6 embryos (CC), 15 from 9 embryos (IDR), 16 from 9 embryos (CC-IDR1), and 10 from 6 embryos. Mean and SE values are shown.

## Discussion

When a molecule demonstrates functions in a cell, it is not always present in the active state; however, its association with regulators often aids regulation of its activity. In some cases, the localization of each molecule is controlled spatially and temporally, and the formation of the complex itself is considered the rate-limiting process, while in other cases, the complex is always formed and external signals induce the activation. The mechanism of cellular regulation of the activity of molecular complexes can be studied comprehensively *in vitro.* To understand the actual cellular regulatory mechanism within a cell, it is necessary to carefully observe the spatiotemporal dynamics of each molecule *in vivo* and to integrate the available knowledge.

Cytoplasmic dynein I is a microtubule-based motor protein that is indispensable for various cellular processes, including the formation, maintenance, and elongation of mitotic spindles (Roberts *et al.*, 2013). Recent *in vitro* studies have revealed the mechanism by which dynein forms complexes with its regulatory proteins and the properties of the complexes (Reck-Peterson *et al.*, 2018; Olenick and Holzbaur, 2019). Here, we focused on the manner in which dynein localized and functioned at mitotic spindles after NEBD and investigated the spatiotemporal dynamics of dynein and its regulatory proteins using *C. elegans* early embryos to understand the mechanism of cellular regulation of dynein.

### Regulatory proteins accumulate earlier than dynein

We found that dynein and its regulatory proteins did not accumulate simultaneously, but accumulation occurred in a sequential manner. Several regulatory proteins, including NUD-2, LIS-1, and LIN-5, gradually accumulated earlier than dynein and dynactin (**Figure 3A**). This accumulation order was not determined by the molecular weight (**Figure 4**).

Among the early accumulating proteins, NUD-2 showed the earliest accumulation that was initiated before CeNEBD (**Figures 3A and 5A**). We hypothesized that the earliest accumulation of NUD-2 contributed to the accumulation of other proteins. In NUD-2-depleted embryos, we found that dynein, dynactin, and LIS-1 gradually accumulated as in the unperturbed embryos, but the protein concentration in the entire spindle region decreased after the initial accumulation (**Figure 7C**). These results indicate that NUD-2 depletion affects the dynamics of later accumulating proteins.

The results obtained via NUD-2 depletion suggest a sequential effect, where earlier accumulation affects the dynamics of the proteins accumulated later. Based on the following considerations, this sequential effect seems to demonstrate implications for the intracellular regulation of cytoplasmic dynein (**Figure 9B**). First, NDEL1, a homolog of NUD-2, recruits LIS1 to dynein (McKenney *et al.*, 2010; Wang *et al.*, 2013). If NUD-2 level decreases in the spindle region, LIS-1 loses one of the interaction partners and leaks out of the spindle region, leading to a reduction in the ratio of dynein bound to LIS-1 in the spindle region. Second, recent studies have shown that LIS1 binding to dynein shifts dynein conformation, promotes dynein-dynactin binding, and dissociates from dynein after the binding of dynactin to dynein (Qiu *et al.*, 2019; Elshenawy *et al.*, 2020; Htet *et al.*, 2020; Marzo *et al.*, 2020); therefore, a decrease in LIS-1 level decreases the proportion of the dynein-dynactin complex. In both types of dynein complexes, the duration of the presence of dynein on microtubules is expected to be longer (McKenney *et al.*, 2010, 2014; Schlager *et al.*, 2014; Elshenawy *et al.*, 2020; Marzo *et al.*, 2020). A decrease in the proportion of dynein with a longer duration on microtubules would decrease the affinity for the entire spindle region, leading to a decrease in the maximum accumulation ratio and the decrease in protein concentration at the spindle region after the initial accumulation.

**Figure 9.**
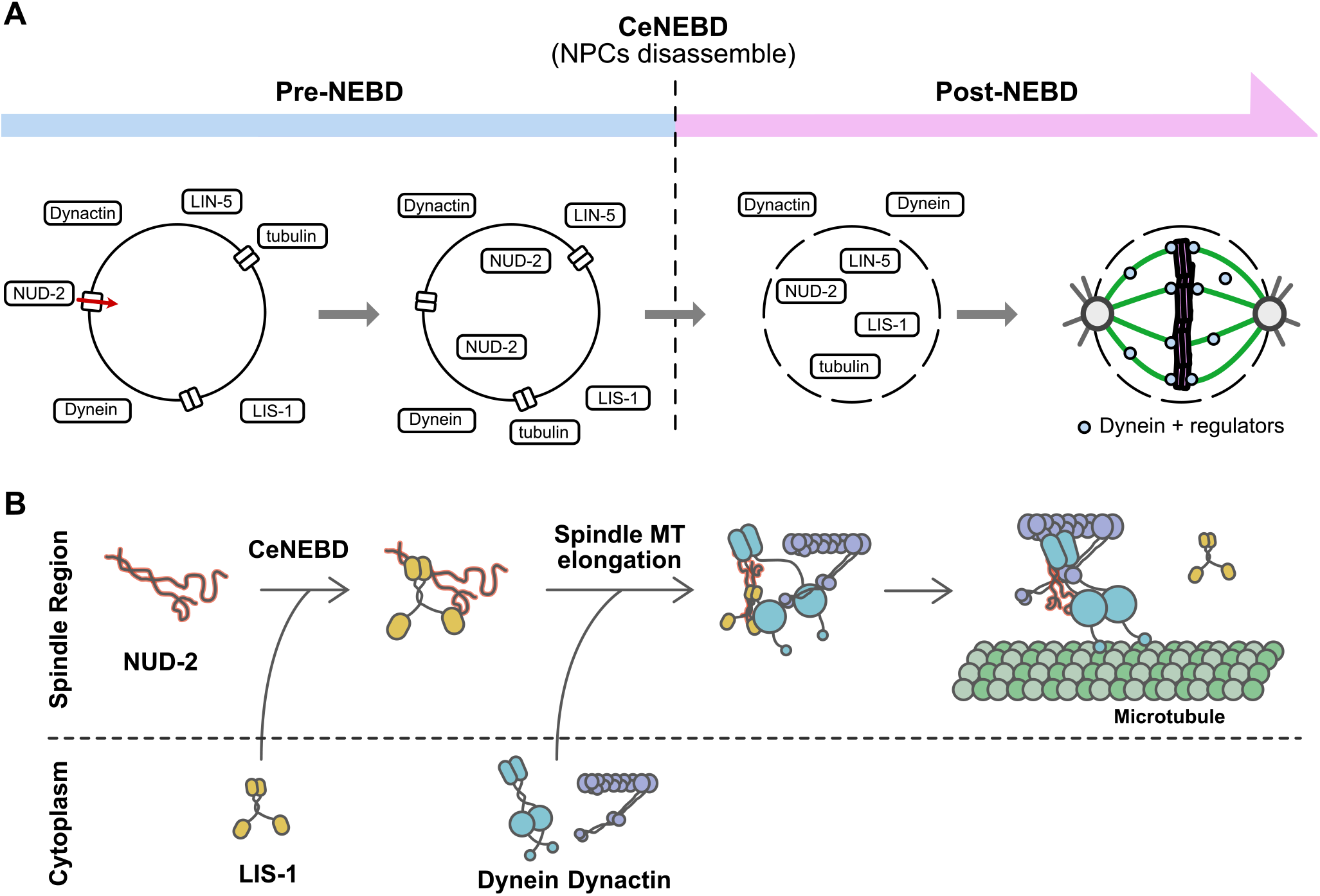
Sequential accumulation of dynein and its regulatory proteins and its implication in dynein regulation. (A) Schematic representation showing the sequential accumulation of the proteins observed in this study. NUD-2 accumulates at the spindle region before CeNEBD. After CeNEBD, LIS-1 and LIN-5 accumulate, followed by dynein and dynactin. Dynein associates with the accumulated proteins and contributes to the formation and maintenance of mitotic spindles. (B) A model of dynein regulation through accumulation. NUD-2, accumulated before CeNEBD, binds to LIS-1, which accumulates after CeNEBD. Following the accumulation of LIS-1, dynein and dynactin accumulate. Dynein then associates with LIS-1, which is supported by NUD-2. The binding of LIS-1 changes dynein conformation and facilitates the complex formation of dynein with dynactin. Upon the binding of dynactin to dynein, LIS-1 dissociates from dynein.

In the sequential regulatory mechanism discussed above, it is important to determine whether the concentrations of the proteins are sufficient to enable binding with each other. Since none of the regulatory proteins observed presented an accumulation level that was markedly lower than that of dynein (**Figure 1D**), it could be implied that the observed proteins were present in the spindle region in sufficient amounts to facilitate binding with dynein.

### Accumulation in the bulk spindle region

We found that the proteins accumulated earlier than dynein and dynactin were first localized at the bulk spindle region (**Figure 2**). This affinity for the bulk spindle region is suggested to enable protein accumulation in the spindle region before the elongation of spindle microtubules. In contrast, dynein and dynactin gradually accumulated through spindle microtubules and localized later to the kinetochores (**Figures 2D and E**). Our previous research showed that DNC-1 accumulated in the nascent spindle region, but not in a uniform manner (Hayashi *et al.*, 2012). Our present study revealed that the accumulation was not in the bulk spindle region, but rather occurred at the kinetochore. The proteins accumulating in the bulk spindle region increase their absolute concentrations in the spindle region through earlier accumulation, resulting in efficient complex formation with dynein.

### Mechanism of protein accumulation in the bulk spindle region

A pertinent aspect worth exploration is the mechanism of accumulation in the bulk spindle region. For accumulation in spindle microtubules and kinetochores, a mechanism through specific protein-protein interactions can be proposed; however, the possible mechanism of accumulation in the bulk space should be elucidated. Considering the microscopic accumulation process, it is hypothesized that the protein molecules may enter the spindle region mainly through diffusion (“on” process) since it occurs after NPCs are disassembled. Some molecules may exit the region through diffusion (“off” process); however, if a mechanism exists for the entrapment of the molecules in the spindle region, it may result in the accumulation of the molecules. The balance between on and off states is assumed to be different depending on the molecule, and the accumulation in the bulk spindle region may be possible when this on-off relationship satisfies certain conditions. The balance between the on and off is also affected by other accumulated proteins. The accumulation of a molecule with a high affinity for the spindle region may increase the affinity of the molecule, which accumulates later. Additionally, it is notable that accumulation was observed to a certain extent, even for dextran, which was not expected to establish interactions specifically with intracellular molecules; such an event was not observed with PEGs (**Figure 4**). The accumulation itself may occur if properties, such as charge, hydrophilicity, and structure, are satisfied. Of course, the possibility that some factors in the nucleus specifically recruit proteins cannot be excluded from this study alone.

### Mechanism and implication of pre-NEBD accumulation of NUD-2

We assumed that the pre-NEBD accumulation of NUD-2 was related to nucleocytoplasmic transport and NUD-2 dynamics in *ran-1* (RNAi) embryos (**Figure 6A**). In RAN-1-depleted embryos, NUD-2 showed only post-NEBD accumulation, suggesting that nucleocytoplasmic transport mediated pre-NEBD accumulation (**Figure 6B**). This RAN-1 independence of post-NEBD accumulation of NUD- 2 was in contrast to that of tubulin (**Figures 6C and 6D**) (Hayashi *et al.*, 2012). Regarding the pre-NEBD accumulation of NUD-2, we also found that the C-terminal putative helix region was essential for the protein injection approach (**Figure 8**). Previous studies have shown that the C-terminal region of NDEL1/NDE1 comprises many phosphorylation sites and a dynein binding site, and can bend back onto the N-terminal coiled-coil (Feng *et al.*, 2000; Niethammer *et al.*, 2000; Sasaki *et al.*, 2000; Yan *et al.*, 2003; Liang *et al.*, 2004; Toyo-Oka *et al.*, 2005; Guo *et al.*, 2006; Mori *et al.*, 2007; Bradshaw *et al.*, 2008; Torisawa *et al.*, 2011; Soares *et al.*, 2012). As suggested by Soares et al., phosphorylation in the C-terminal region might affect the overall molecular structure of NUD-2 and could induce pre-NEBD accumulation.

### Applications of protein injection approach

Previous studies have reported the use of exogenous polymers to investigate the permeability of nuclear membranes (Galy *et al.*, 2003; Updike *et al.*, 2011), and we used the same technique in this study (**Figure 4**). In this study, we have shown that the intracellular dynamics of recombinant proteins can be observed by injecting proteins purified from *E. coli* (**Figure 8**). The advantage of this method is that the results can be obtained in a shorter time (typically in 1 week or 2 weeks) than the observations obtained using transgenic worms, which requires the establishment of new worm strains.

### Universality and diversity of accumulation and its role in cellular regulation of dynein

Nuclear accumulation of proteins during mitosis has also been observed in *D. melanogaster* (Yao *et al.*, 2012; Schweizer *et al.*, 2015; Métivier *et al.*, 2021). One characteristic shared by both organisms is that their mitosis is semi-open (Makarova and Oliferenko, 2016), in which the nuclear envelope disrupts only partially, not completely, during mitosis. The affinity of the substance for the spindle region of the bulk may differ between cases in which all NEs collapse (open mitosis) and those in which they do not (semi-open or semi-closed).

The accumulation of components required for spindle formation is essential for mitosis, and dynein is a crucial molecule in this process. It will be desirable to study the spatiotemporal dynamics of dynein and its regulatory proteins in species with different modes of mitosis and to compare the mechanisms employed in individual organisms to achieve the universality and diversity of the accumulation phenomenon.

## MATERIALS & METHODS

### *C. elegans* strains

The worm strains used in this study have been summarized in Table S1 and were maintained at 22 °C on standard nematode growth medium (NGM) plates with OP50 *Escherichia coli.* To establish worm strains expressing both histone and dynein-regulatory proteins, we used LP451 (expressing NUD- 2::mNG), LP563 (mNG::DNC-1), LP585 (LIN-5::mNG), and LP591 (LIS-1::mNG), provided by the *Caenorhabditis* Genetics Center. The strains were subjected to crossing experiments with CAL0941 that expressed mCherry-fused HIS-58. CAL2221, which was used to visualize dynein, was established for this study using the CRISPR/Cas9 method (Dickinson and Goldstein, 2016). The guide RNA was designed to the 5’-terminal of exon1 of *dhc-1* and the rescued fragment containing full-length *dhc-1* sequence fused with hsGFP was injected into the gonads of young adult N2 worms with pRF4, a plasmid used for *rol* mutant screening. hsGFP is a recombinant GFP containing 6xHis- tag and streptavidin-binding peptide tag (Kobayashi *et al.*, 2008). After performing screening of the *rol* mutant, the worms were screened using fluorescence signals. After confirming the insertion via sequencing, CAL2221 was subjected to crossing experiments with CAL0941 to visualize dynein and histones simultaneously.

### RNAi experiments

For the synthesis of double-stranded RNAs (dsRNAs), oligonucleotides containing T3 and T7 promoters were used. The sequences of the oligonucleotides were the same as those available in PhenoBank (https://worm.mpi-cbg.de/phenobank/cgi-bin/ProjectInfoPage.py). The dsRNA sequences were amplified from the genomic DNA of the N2 strain. After amplification, the dsRNAs were synthesized from the products using T3 and T7 RNA polymerases (Promega, P2075, and P2083). The transcription products were incubated at 70 °C for 10 min and at 37 °C for 30 min for annealing. After annealing, the products were filtered using SUPREC™-01 (Takara, 9040). To inject the purified dsRNAs, young adult worms were placed on a thin layer of 2% agarose (Lonza, SeaKem LE agarose) on a 24 × 55-mm coverslip (Matsunami). After covering the worms with halocarbon oil (Sigma, H8898-50MK), the coverslip was mounted and analyzed using an inverted microscope (Axiovert 100, Carl Zeiss). The dsRNAs were injected into the worms using a microinjector (Eppendorf, FemtoJet). After the completion of injection, 5–10 μL of M9 buffer (22 mM KH2PO4, 42 mM Na2HPO4, and 86 mM NaCl) was added to the oil to recover the worms. The worms were transferred to a new NGM plate with OP50 *E. coli* and were incubated at 22 °C for 44–48 h (*nud-2*), 24-28 h (*tbb-2*), or 16–20 h (*ran-1* and *cdk-1*) before conducting observations.

### Construction and purification of recombinant proteins

The full-length coding sequence of *nud-2* was amplified from the cDNA of the N2 strain using the KOD One PCR Master Mix (Toyobo, KMM-101). The amplified sequence was inserted into the pET17b vector (Invitrogen) together with the sequence of SBP-mCherry using seamless cloning with the NEBuilder HiFi DNA Assembly Master Mix (New England BioLabs, E2621). To construct the truncated fragments, unnecessary sequences were removed from the full-length constructs using seamless cloning. The plasmids were transformed into Rosetta2 (DE3) competent cells (Novagen, 71397). The recombinant proteins were purified using SBP-tag and StrepTactin Sepharose. *E. coli* cells obtained from 500 mL culture were harvested using centrifugation for 10 min at 4800 rpm (Beckman, Allegra-30XR), and were subjected to freezing in liquid nitrogen. The collected cells were suspended in lysis buffer (50 mM HEPES-KOH, 150 mM NaCl, 1 mM EGTA, 10% (w/v) sucrose, and pH7.2) supplemented with the ProteoGuard EDTA-free protease inhibitor cocktail (Clontech, 635673). The cells in the suspended solution were sonicated using the Q125 sonicator (Qsonica) and the following settings: 60% amplitude, +4 °C water bath, and 1-s ON/1-s OFF pulses. The total sonication time was 10 min. The homogenized solution was centrifuged at 75,000 rpm for 15 min (Beckman, TL100.3). The supernatant was loaded onto a StrepTactin Sepharose column with a volume of 1 mL, followed by washing with lysis buffer. The proteins were eluted with lysis buffer supplemented with 2.5 mM desthiobiotin. Protein concentrations were determined via the Bradford method using the TaKaRa Bradford Protein Assay Kit (Takara Bio, T9310A).

### Gonad injection of recombinant proteins or polymers

Purified proteins or polymers were diluted using 1× PBS (Takara Bio, T900), and loaded into custom-made microneedles prepared with the P1000IVF micropipette puller (Sutter Instrument). Young adult worms were placed on a thin layer of 2% agarose (Lonza, SeaKem LE agarose) prepared on a 24 × 55-mm coverslip (Matsunami). After covering the worms with halocarbon oil, the coverslips were mounted and analyzed using the Axiovert100 inverted microscope (Carl Zeiss). Protein solutions were injected into the worms using the FemtoJet microinjector (Eppendorf). After the completion of injection, 5–10 μL of M9 buffer was added to release the worms. After the release, the worms were transferred to a new NGM plate and were incubated at 22 °C for at least 3 h for incorporation of the injected components into the embryos.

### Imaging of *C. elegans* early embryos

Worms were dissected using a 0.75 × egg salt buffer (118mM NaCl, 40mM KCl, 3.4mM CaCl2, 3.4mM MgCl2, 5mM HEPES pH 7.2). The embryos from the dissected worms were mounted in 0.75 × egg salt buffer, which was placed on a 26 × 76-mm custom-made coverslip (Matsunami). To eliminate the effects of deformation, no coverslip was mounted on the embryos. Egg-mounted coverslips were set to a spinning-disk confocal fluorescent microscope consisting of the IX71 inverted microscope (Olympus) and the CSU-X1 spinning-head (Yokogawa). The microscope was equipped with a 60× silicon-immersion objective lens (Olympus, UPlanSApo, 60x/1.30Sil) and a 2.0× intermediate magnification lens. Images were acquired using an EM-CCD (Andor, iXon) managed by the NIS elements software (Nikon) at 10-s intervals. In single-slice acquisition, the exposure time was 180 ms for both the 488-nm and 561-nm channels. To acquire 3D images, a piezo-actuated microscope stage (PI) was used and the acquisition interval was set to 20 s. In the 3D observations, the exposure times were 120 ms for the 488-nm channel and 60 ms for the 561-nm channel. In the observations of nocodazole-treated embryos, worms were dissected using a 0.75 × egg salt buffer supplemented with 10 μg/mL nocodazole (Fujifilm Wako, 140-08531). After dissection, the worms were mounted using the same buffer. The experimental room was air-conditioned and temperature was maintained at 21–23°C.

### Image analysis

To analyze the images of *C. elegans* early embryos, the background intensity was subtracted, and photobleaching effect was corrected by assuming an exponential model. After preprocessing, the mean CI and mean NI values were measured to calculate the N/C ratio of the intensity. Mean CI was calculated by averaging the mean intensities measured in three circular regions with a 100-pixel diameter (13.6 μm) randomly placed in the early 1-cell stage embryos, which were in the stage before the growth of pronuclei. We selected this timing because all proteins observed showed relatively uniform subcellular distributions in the whole embryo. The calculated mean CI was used to determine the N/C ratio throughout the calculations. In the measurements of NIs, we manually selected the nuclear boundary using the signals of histones or accumulated proteins. In the measurement of LIN-5 and tubulin, which presented with strong signals at centrosomes, centrosomal signals were masked with a circular region with a 30-pixel diameter (4.09 μm).

### Statistical analyses

All graphs were generated, and statistical analyses were performed using Prism v.7 (GraphPad). For all figures, the error bars represent the standard error of the mean. For all experiments, data were obtained from independent experiments using embryos that were independent of the worms.

**Table 1.** Quantification of the accumulation of dynein and the regulatory proteins

|  | **Cytoplasmic** | **Nuclear** | **Maximum N/C** | **n** |
| --- | --- | --- | --- | --- |
|  | **intensity (CI)** | **intensity (NI)** | **ratio** |  |
| | $\times 10^2$ a. u. | $\times 10^2$ a. u. | | |
| **NUD-2** | 1.4 ± 0.2 | 6.3 ± 1.9 | 4.6 ± 0.6 | 8 |
| **LIS-1** | 9.0 ± 1.7 | 13.1 ± 2.6 | 1.5 ± 0.1 | 11 |
| **LIN-5** | 5.9 ± 1.1 | 10.2 ± 2.3 | 1.7 ± 0.2 | 10 |
| **DNC-1** | 3.4 ± 0.7 | 5.7 ± 1.8 | 1.6 ± 0.2 | 11 |
| **DHC-1** | 2.1 ± 0.6 | 3.7 ± 1.2 | 1.7 ± 0.1 | 9 |

## ACKNOWLEDGMENTS

Some strains were provided by the *Caenorhabditis* Genetics Center funded by the NIH Office of Research Infrastructure Programs (P40 OD010440). We would like to thank Dr. Kei Saito (National Institute of Genetics) for reading the manuscript and for providing helpful comments. This work was supported by JSPS KAKENHI (grant numbers JP19K16094 to TT, JP18H02414 to AK, and JP18KK0202 to AK and KK).

## REFERENCES

Aguirre-Chen C, Bülow HE, Kaprielian Z (2011). C. elegans bicd-1, homolog of the Drosophila dynein accessory factor Bicaudal D, regulates the branching of PVD sensory neuron dendrites. Development 138, 507–518.

Askjaer P, Galy V, Hannak E, Mattaj IW (2002). Ran GTPase cycle and importins alpha and beta are essential for spindle formation and nuclear envelope assembly in living Caenorhabditis elegans embryos. Mol Biol Cell 13, 4355–4370.

Aumais JP, Williams SN, Luo W, Nishino M, Caldwell KA, Caldwell GA, Lin S-H, Yu-Lee, L-Y (2003). Role for NudC, a dynein-associated nuclear movement protein, in mitosis and cytokinesis. J Cell Sci 116, 1991–2003.

Bader JR, Vaughan KT (2010). Dynein at the kinetochore: timing, interactions, and functions. Semin Cell Dev Biol 21, 269–275.

Bamba C, Bobinnec Y, Fukuda M, Nishida E (2002). The GTPase Ran regulates chromosome positioning and nuclear envelope assembly in vivo. Curr Biol 12, 503–507.

Bradshaw NJ, Ogawa F, Antolin-Fontes B, Chubb JE, Carlyle BC, Christie S, Claessens A, Porteous DJ, Millar JK (2008). DISC1, PDE4B, and NDE1 at the centrosome and synapse. Biochem Biophys Res Commun 377, 1091–1096.

Cockell MM, Baumer K, Gönczy P (2004). lis-1 is required for dynein-dependent cell division processes in C. elegans embryos. J Cell Sci 117, 4571–4582.

Cross RA, McAinsh A (2014). Prime movers: the mechanochemistry of mitotic kinesins. Nat Rev Mol Cell Biol 15, 257–271.

Derewe U et al. (2007). The structure of the coiled-coil domain of Ndel1 and the basis of its interaction with Lis1, the causal protein of Miller-Dieker lissencephaly. Structure 15, 1467–1481.

Dickinson DJ, Goldstein B (2016). CRISPR-based methods for Caenorhabditis elegans genome engineering. Genetics 202, 885–901.

Elshenawy MM, Kusakci E, Volz S, Baumbach J, Bullock SL, Yildiz A (2020). Lis1 activates dynein motility by modulating its pairing with dynactin. Nat Cell Biol 22, 570–578.

Feng Y, Olson EC, Stukenberg PT, Flanagan LA, Kirschner MW, Walsh, CA (2000). LIS1 regulates CNS lamination by interacting with mNudE, a central component of the centrosome. Neuron 28, 665–679.

Galy V, Mattaj IW, Askjaer P (2003). Caenorhabditis elegans nucleoporins Nup93 and Nup205 determine the limit of nuclear pore complex size exclusion in vivo. Mol Biol Cell 14, 5104–5115.

Gassmann R et al. (2008). A new mechanism controlling kinetochore-microtubule interactions revealed by comparison of two dynein-targeting components: SPDL-1 and the Rod/Zwilch/Zw10 complex. Genes Dev 22, 2385–2399.

Gönczy P et al. (2000). Functional genomic analysis of cell division in C. elegans using RNAi of genes on chromosome III. Nature 408, 331–336.

Guo J, Yang Z, Song W, Chen Q, Wang F, Zhang Q, Zhu X (2006). Nudel contributes to microtubule anchoring at the mother centriole and is involved in both dynein-dependent and - independent centrosomal protein assembly. Mol Biol Cell 17, 680–689.

Hachet V, Canard C, Gönczy P (2007). Centrosomes promote timely mitotic entry in C. elegans embryos. Dev Cell 12, 531–541.

Hayashi H, Kimura K, Kimura A (2012). Localized accumulation of tubulin during semi-open mitosis in the Caenorhabditis elegans embryo. Mol Biol Cell 23, 1688–1699.

Heald R, Khodjakov A (2015). Thirty years of search and capture: the complex simplicity of mitotic spindle assembly. J Cell Biol 211, 1103–1111.

Heppert JK, Dickinson DJ, Pani AM, Higgins CD, Steward A, Ahringer J, Kuhn JR, Goldstein B (2016). Comparative assessment of fluorescent proteins for in vivo imaging in an animal model system. Mol Biol Cell 27, 3385–3394.

Heppert JK, Pani AM, Roberts AM, Dickinson DJ, Goldstein, B (2018). A CRISPR tagging-based screen reveals localized players in Wnt-directed asymmetric cell division. Genetics 208, 1147–1164.

Hirokawa N, Noda Y, Tan Y, Niwa S (2009). Kinesin superfamily motor proteins and intracellular transport. Nat Rev Mol Cell Biol 10, 682–696.

Htet ZM, Gillies JP, Baker RW, Leschziner AE, DeSantis ME, Reck-Peterson SL (2020). LIS1 promotes the formation of activated cytoplasmic dynein-1 complexes. Nat Cell Biol 22, 518–525.

Kardon JR, Vale RD (2009). Regulators of the cytoplasmic dynein motor. Nat Rev Mol Cell Biol 10, 854–865.

King SM (2011). Dyneins: Structure, Biology and Disease, Academic Press.

Kiyomitsu T (2019). The cortical force-generating machinery: how cortical spindle-pulling forces are generated. Curr Opin Cell Biol 60, 1–8.

Kobayashi T, Morone N, Kashiyama T, Oyamada H, Kurebayashi N, Murayama T (2008). Engineering a novel multifunctional green fluorescent protein tag for a wide variety of protein research. PLoS One 3, e3822.

Lee KK, Gruenbaum Y, Spann P, Liu J, Wilson KL (2000). C. elegans nuclear envelope proteins emerin, MAN1, lamin, and nucleoporins reveal unique timing of nuclear envelope breakdown during mitosis. Mol Biol Cell 11, 3089–3099.

Liang Y, Yu W, Li Y, Yang Z, Yan X, Huang Q, Zhu X (2004). Nudel functions in membrane traffic mainly through association with Lis1 and cytoplasmic dynein. J Cell Biol 164, 557–566.

Makarova M, Oliferenko S (2016). Mixing and matching nuclear envelope remodeling and spindle assembly strategies in the evolution of mitosis. Curr Opin Cell Biol 41, 43–50.

Malone CJ, Misner L, Le Bot N, Tsai M-C, Campbell JM, Ahringer J, White JG (2003). The C. elegans hook protein, ZYG-12, mediates the essential attachment between the centrosome and nucleus. Cell 115, 825–836.

Marzo MG, Griswold JM, Markus SM (2020). Pac1/LIS1 stabilizes an uninhibited conformation of dynein to coordinate its localization and activity. Nat Cell Biol 22, 559–569.

McKenney RJ, Huynh W, Tanenbaum ME, Bhabha G, Vale RD (2014). Activation of cytoplasmic dynein motility by dynactin-cargo adapter complexes. Science 345, 337–341.

McKenney RJ, Vershinin M, Kunwar A, Vallee RB, Gross SP (2010). LIS1 and NudE induce a persistent dynein force-producing state. Cell 141, 304–314.

Métivier M et al. (2021). Drosophila tubulin-specific chaperone E recruits tubulin around chromatin to promote mitotic spindle assembly. Curr Biol 31, 684–695.e6.

Mori D et al. (2007). NDEL1 phosphorylation by Aurora-A kinase is essential for centrosomal maturation, separation, and TACC3 recruitment. Mol Cell Biol 27, 352–367.

Niethammer M, Smith DS, Ayala R, Peng J, Ko J, Lee MS, Morabito M, Tsai LH (2000). NUDEL is a novel Cdk5 substrate that associates with LIS1 and cytoplasmic dynein. Neuron 28, 697–711.

Olenick MA, Holzbaur ELF (2019). Dynein activators and adaptors at a glance. J Cell Sci 132.

Pfister KK, Shah PR, Hummerich H, Russ A, Cotton J, Annuar AA, King SM, Fisher EMC (2006). Genetic analysis of the cytoplasmic dynein subunit families. PLoS Genet 2, e1.

Portegijs V, Fielmich L-E, Galli M, Schmidt R, Muñoz J, van Mourik T, Akhmanova A, Heck AJR, Boxem M, van den Heuvel S (2016). Multisite phosphorylation of NuMA-related LIN-5 controls mitotic spindle positioning in C. elegans. PLoS Genet 12, e1006291.

Portier N, Audhya A, Maddox PS, Green RA, Dammermann A, Desai A, Oegema K (2007). A microtubule-independent role for centrosomes and aurora a in nuclear envelope breakdown. Dev Cell 12, 515–529.

Qiu R, Zhang J, Xiang X (2019). LIS1 regulates cargo-adapter-mediated activation of dynein by overcoming its autoinhibition in vivo. J Cell Biol 218, 3630–3646.

Raaijmakers JA, Medema RH (2014). Function and regulation of dynein in mitotic chromosome segregation. Chromosoma 123, 407–422.

Raaijmakers JA, Tanenbaum ME, Medema RH (2013). Systematic dissection of dynein regulators in mitosis. J Cell Biol 201, 201–215.

Reck-Peterson SL, Redwine WB, Vale RD, Carter AP (2018). The cytoplasmic dynein transport machinery and its many cargoes. Nat Rev Mol Cell Biol 19, 382–398.

Roberts AJ, Kon T, Knight PJ, Sutoh K, Burgess SA (2013). Functions and mechanics of dynein motor proteins. Nat Rev Mol Cell Biol 14, 713–726.

Sasaki S, Shionoya A, Ishida M, Gambello MJ, Yingling J, Wynshaw-Boris A, Hirotsune S (2000). A LIS1/NUDEL/cytoplasmic dynein heavy chain complex in the developing and adult nervous system. Neuron 28, 681–696.

Schlager MA, Hoang HT, Urnavicius L, Bullock SL, Carter AP (2014). In vitro reconstitution of a highly processive recombinant human dynein complex. EMBO J 33, 1855–1868.

Schmidt R, Fielmich L-E, Grigoriev I, Katrukha EA, Akhmanova A, van den Heuvel S (2017). Two populations of cytoplasmic dynein contribute to spindle positioning in C. elegans embryos. J Cell Biol 216, 2777–2793.

Schweizer N, Pawar N, Weiss M, Maiato H (2015). An organelle-exclusion envelope assists mitosis and underlies distinct molecular crowding in the spindle region. J Cell Biol 210, 695–704.

Simões PA, Celestino R, Carvalho AX, Gassmann R (2018). NudE regulates dynein at kinetochores but is dispensable for other dynein functions in the C. elegans early embryo. J Cell Sci 131, jcs212159.

Soares DC et al. (2012). The mitosis and neurodevelopment proteins NDE1 and NDEL1 form dimers, tetramers, and polymers with a folded back structure in solution. J Biol Chem 287, 32381–32393.

Torisawa T, Ichikawa M, Furuta A, Saito K, Oiwa K, Kojima H, Toyoshima YY, Furuta K (2014). Autoinhibition and cooperative activation mechanisms of cytoplasmic dynein. Nat Cell Biol 16, 1118–1124.

Torisawa T, Kimura A (2020). The Generation of Dynein Networks by Multi-Layered Regulation and Their Implication in Cell Division. Front Cell Dev Biol 8, 22.

Torisawa T, Nakayama A, Furuta K ‘ya, Yamada M, Hirotsune S, Toyoshima YY (2011). Functional dissection of LIS1 and NDEL1 towards understanding the molecular mechanisms of cytoplasmic dynein regulation. J Biol Chem 286, 1959–1965.

Toya M, Terasawa M, Nagata K, Iida Y, Sugimoto A (2011). A kinase-independent role for Aurora A in the assembly of mitotic spindle microtubules in Caenorhabditis elegans embryos. Nat Cell Biol 13, 708–714.

Toyo-Oka K et al. (2005). Recruitment of katanin p60 by phosphorylated NDEL1, an LIS1 interacting protein, is essential for mitotic cell division and neuronal migration. Hum Mol Genet 14, 3113–3128.

Tzur YB, Gruenbaum Y (22000-2013). Nuclear envelope breakdown and reassembly in C. elegans: evolutionary aspects of lamina structure and function. In: Madame Curie Bioscience Database [Internet]. Austin (TX): Landes Bioscience. Available from: https://www.ncbi.nlm.nih.gov/books/NBK6297/

Updike DL, Hachey SJ, Kreher J, Strome S (2011). P granules extend the nuclear pore complex environment in the C. elegans germ line. J Cell Biol 192, 939–948.

Vaisberg EA, Koonce MP, McIntosh JR (1993). Cytoplasmic dynein plays a role in mammalian mitotic spindle formation. J Cell Biol 123, 849–858.

Wang S, Ketcham SA, Schön A, Goodman B, Wang Y, Yates J, 3rd, Freire E, Schroer TA, Zheng Y (2013). Nudel/NudE and Lis1 promote dynein and dynactin interaction in the context of spindle morphogenesis. Mol Biol Cell 24, 3522–3533.

Yamada M, Toba S, Yoshida Y, Haratani K, Mori D, Yano Y, Mimori-Kiyosue Y, Nakamura T, Itoh K, Fushiki S, Setou M, Wynshaw-Boris A, Torisawa T, Toyoshima YY, Hirotsune S (2008). LIS1 and NDEL1 coordinate the plus-end-directed transport of cytoplasmic dynein. EMBO J 27, 2471–2483.

Yan X, Li F, Liang Y, Shen Y, Zhao X, Huang Q, Zhu X (2003). Human Nudel and NudE as regulators of cytoplasmic dynein in poleward protein transport along the mitotic spindle. Mol Cell Biol 23, 1239–1250.

Yao C, Rath U, Maiato H, Sharp D, Girton J, Johansen KM, Johansen, J (2012). A nuclear-derived proteinaceous matrix embeds the microtubule spindle apparatus during mitosis. Mol Biol Cell 23, 3532–3541.

Zhang K, Foster HE, Rondelet A, Lacey SE, Bahi-Buisson N, Bird AW, Carter AP (2017). Cryo-EM reveals how human cytoplasmic dynein is auto-inhibited and activated. Cell 169, 1303–1314.e18.

Zyłkiewicz E, Kijańska M, Choi W-C, Derewenda U, Derewenda ZS, Stukenberg PT (2011). The N-terminal coiled-coil of Ndel1 is a regulated scaffold that recruits LIS1 to dynein. J Cell Biol 192, 433–445.

