## Supplemental Text for "Sequential accumulation of dynein and its regulatory proteins at the spindle region in the *Caenorhabditis elegans* embryo"

### Supplementary Figures

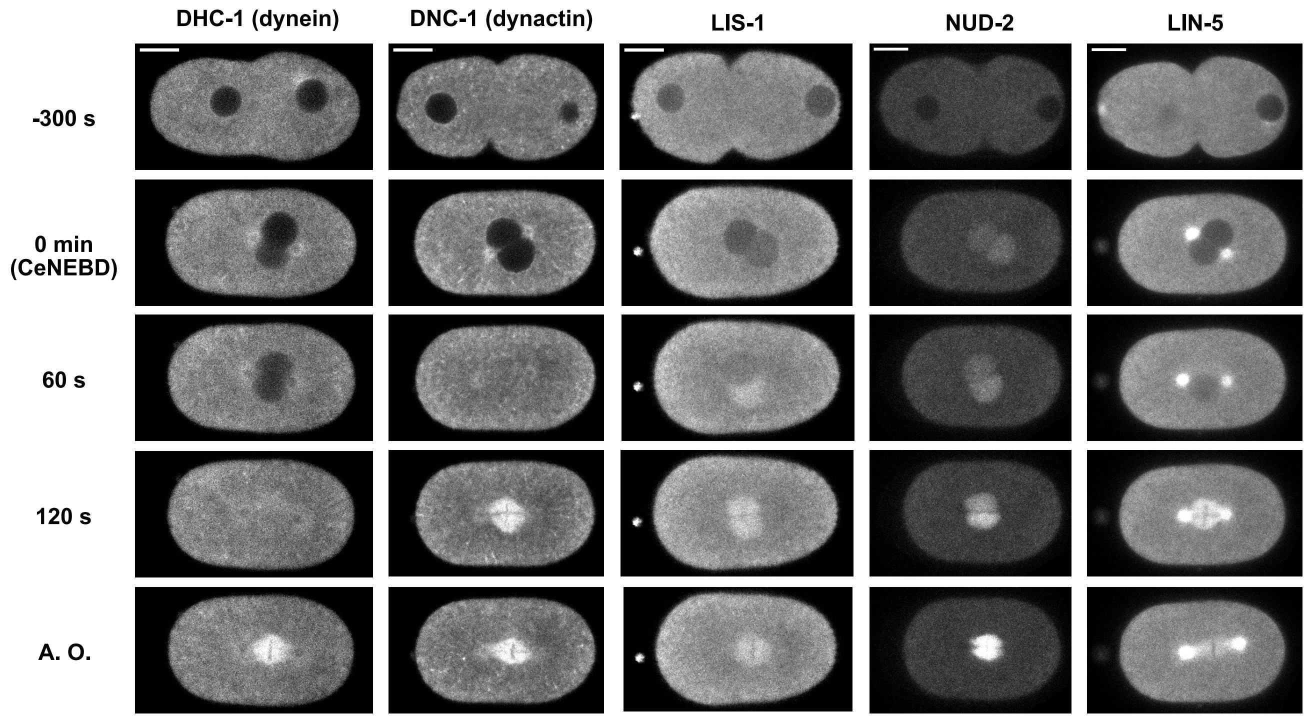


**Figure S1.** Accumulation of dynein, dynactin, LIS-1, NUD-2, and LIN-5.
Typical single-plane time-lapse images showing the temporal dynamics of the proteins indicated above. The left side of the image corresponds to the anterior. The indicated times are relative to CeNEBD. AO denotes anaphase onset. The scale bars indicate 10 µm.

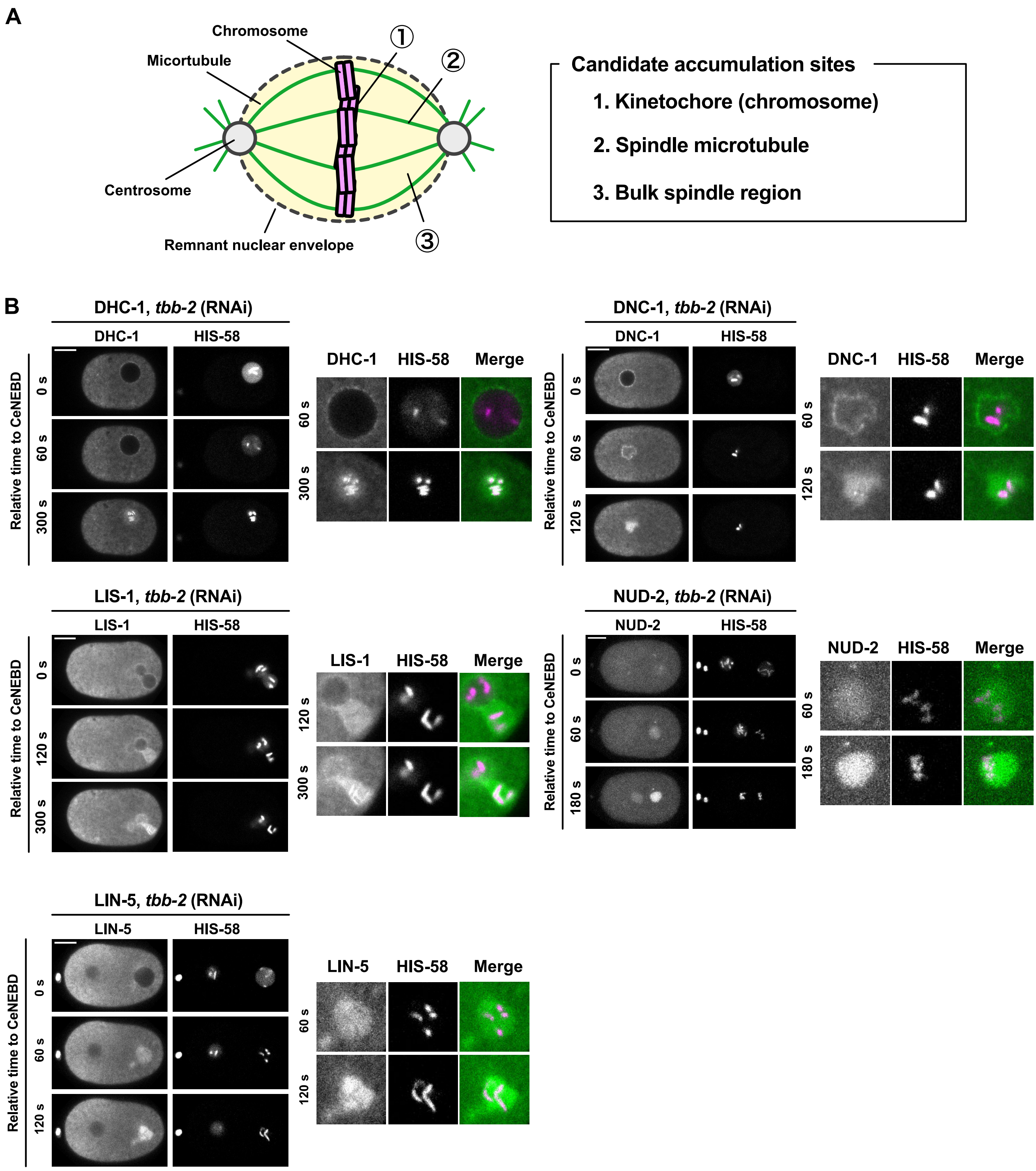


**Figure S2.** Variations in accumulation sites of dynein and its regulatory proteins.

(A) Schematic representation of accumulation site candidates. (B) Accumulation patterns in the tbb-2 embryos. The spatial distribution of LIS-1, NUD-2, LIN-5, dynein, and dynactin in *tbb-2 (RNAi)* embryos is presented. Single-plane time-lapse images of whole embryos and a magnified image of the male pronuclei are shown. The left side of the image corresponds to the anterior. The scale bars indicate 10 μm.

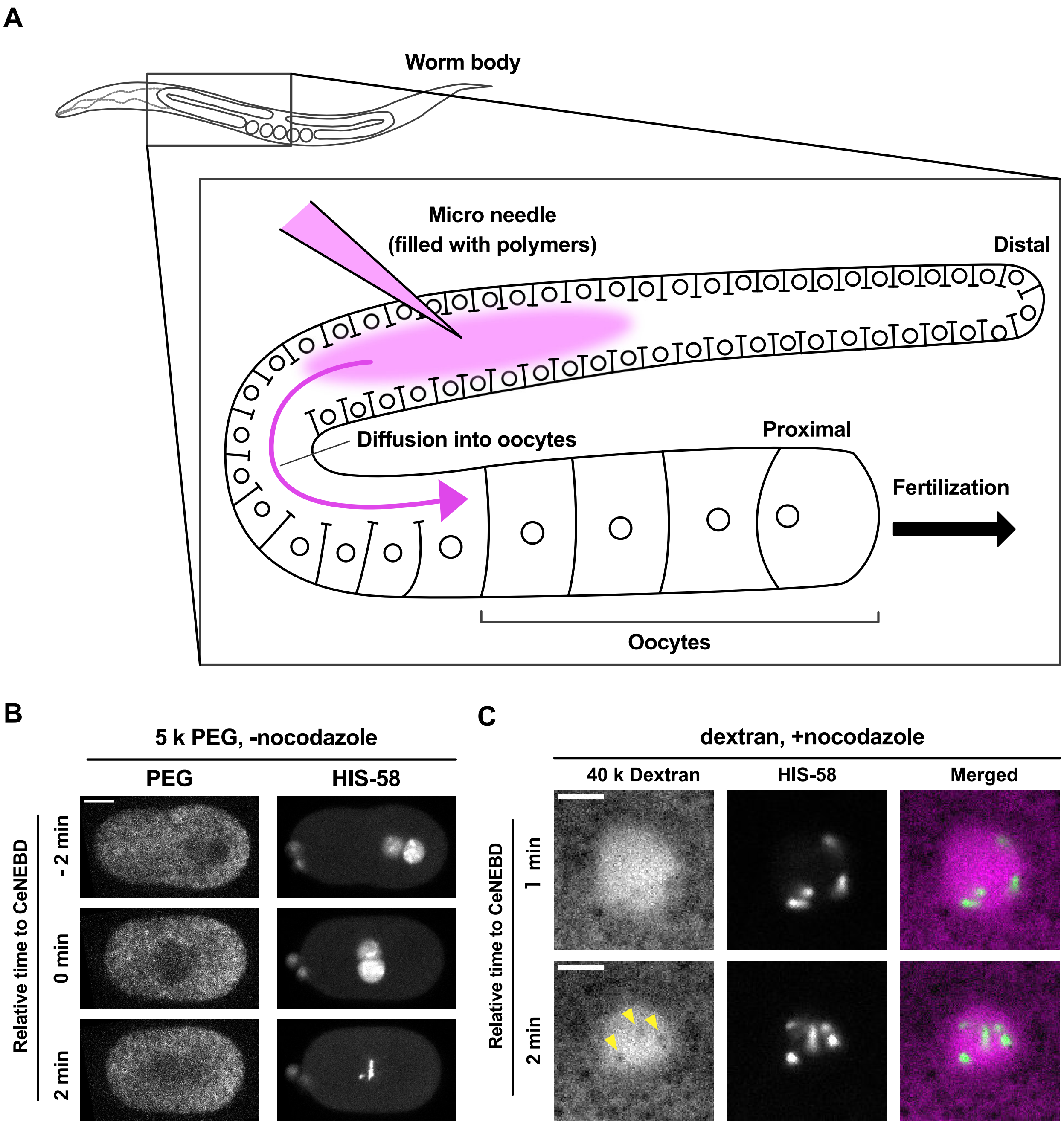


**Figure S3**. Observation of polymer incorporated through gonad injection. (A) Schematic representation of polymer injection experiments. (B) Typical single-plane time-lapse images showing the accumulation pattern of PEG (5 k). The left side of the image corresponds to the anterior. (C) Magnified images showing the distribution of dextran (40 k) in the female pronucleus. The yellow arrowheads indicate the lack of dextran signal in the sites of histone signals. The scale bars indicate 10 μm in (B) and 5 μm in (C).

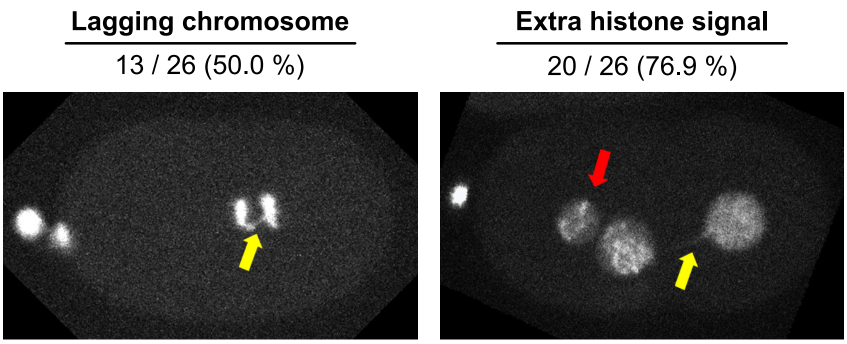


**Figure S4**. Abnormalities in chromosome dynamics in NUD-2 depleted embryos.

The images depict the histone signals in *C. elegans* embryos at approximately first mitotic division. The yellow and red arrows indicate the lagging chromosomes and additional histone signals, respectively. The left side of the image corresponds to the anterior region.

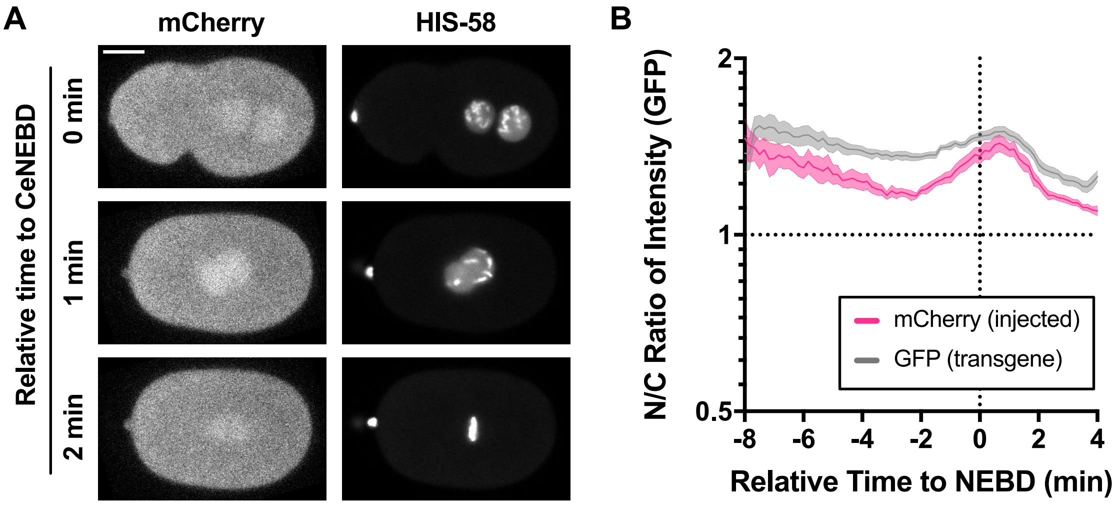


**Figure S5.** Temporal dynamics of injected mCherry. (A) Typical single-plane time-lapse images showing the temporal dynamics of SBP-mCherry incorporated into the early embryo. The right side of the image corresponds to the anterior. The scale bar indicates 10 μm. (B) Time series of the N/C ratio of SBP-mCherry. For comparison, the time series of the N/C ratio of transgenic GFP are also shown. Mean and SE are shown.

**Table S1.** The worm strains used in this study

| | Strain | Genotype | Comment | Figure | Source | Reference |
|---|---|---|---|---|---|---|
| **1** | WH223 | ojIs9 [zyg-12(all)::GFP + unc-119(+)]. | | Figure 1 | CGC | Malone, 2003 |
| **2** | SV1619 | dhc-1(he250[mCherry::dhc-1]) I. | | - | CGC | Schmidt, 2017 |
| **3** | LP439 | nud-2(cp170[nud-2::mNG-C1^3xFlag]) I. | | - | CGC | Heppert, 2018 |
| **4** | LP451 | bicd-1(cp180[mNG-C1^3xFlag::bicd-1]) IV. | | Figure 1 | CGC | Heppert, 2018 |
| **5** | LP563 | dnc-1(cp271[dnc::mNG-C1^3xFlag]) IV. | | - | CGC | Heppert, 2018 |
| **6** | LP585 | lin-5(cp288[lin-5::mNG-C1^3xFlag]) II. | | - | CGC | Heppert, 2018 |
| **7** | LP591 | lis-1(cp294[lis-1::mNG-C1^3xFlag]) III. | | - | CGC | Heppert, 2018 |
| **8** | CAL0234 | ruIs32 [pie-1p::GFP::H2B + unc-119(+)] III. | | Figure 3<br>Figure 8 | This work | |
| **9** | CAL0491 | unc-119(ed3) III; ltIs37[pAA64; pie-1::mCherry::HIS-58; unc-119(+)]; ruIs57[unc-119(+); pie-1p::gfp::tubulin] | | Figure 3<br>Figure 6 | This work | |
| **10** | CAL941 | unc-119 (ed3); wjIs108[unc-119: pie-1 5' : mCherry-his-58: pie-1 3'] | | - | This work | |
| **11** | CAL2221 | dhc-1(hsGFP::dhc-1) I. | | - | This work | |
| **12** | CAL2261 | dnc-1(cp271[dnc::mNG-C1^3xFlag]) IV. : dhc-1(he250[mCherry::dhc-1]) I. | SV1619 x LP563 | Figure 3 | This work | |
| **13** | CAL2271 | lis-1(cp294[lis-1::mNG-C1^3xFlag]) III. : dhc-1(he250[mCherry::dhc-1]) I. | SV1619 x LP591 | Figure 3 | This work | |
| **14** | CAL2281 | lin-5(cp288[lin-5::mNG-C1^3xFlag]) II. : dhc-1(he250[mCherry::dhc-1]) I. | SV1619 x LP585 | Figure 3 | This work | |
| **15** | CAL2291 | nud-2(cp170[nud-2::mNG-C1^3xFlag]) I.: dhc-1(he250[mCherry::dhc-1]) I. | SV1619 x LP439 | Figure 3 | This work | |
| **16** | CAL2302 | nud-2(cp170[nud-2::mNG-C1^3xFlag]) I.: unc-119 (ed3); wjIs108[unc-119: pie-1 5' : mCherry-his-58: pie-1 3'] | LP439 x CAL941 | Figure 1<br>Figure 2<br>Figure 3<br>Figure 5<br>Figure 6 | This work | |
| **17** | CAL2311 | dnc-1(cp271[dnc::mNG-C1^3xFlag]) IV. : unc-119 (ed3); wjIs108[unc-119: pie-1 5' : mCherry-his-58: pie-1 3'] | LP563 x CAL941 | Figure 1<br>Figure 2<br>Figure 3<br>Figure 7 | This work | |
| **18** | CAL2331 | lin-5(cp288[lin-5::mNG-C1^3xFlag]) II. : unc-119 (ed3); wjIs108[unc-119: pie-1 5' : mCherry-his-58: pie-1 3'] | LP585 x CAL941 | Figure 1<br>Figure 2<br>Figure 3<br>Figure 7 | This work | |
| **19** | CAL2341 | lis-1(cp294[lis-1::mNG-C1^3xFlag]) III. : unc-119 (ed3); wjIs108[unc-119: pie-1 5' : mCherry-his-58: pie-1 3'] | LP591 x CAL941 | Figure 1<br>Figure 2<br>Figure 3<br>Figure 7 | This work | |
| **20** | CAL2391 | dhc-1(hsGFP::dhc-1) I. : unc-119 (ed3); wjIs108[unc-119: pie-1 5' : mCherry-his-58: pie-1 3'] | CAL2221 x CAL941 | Figure 1<br>Figure 2<br>Figure 3<br>Figure 7 | This work | |

#### Supplementary Movies

**Movie S1.** Accumulation dynamics of dynein, dynactin, LIS-1, NUD-2, and LIN-5. 100 × Real-Time. Scale bar: 10 µm.

**Movie S2.** Accumulation dynamics of dynein, dynactin, LIS-1, NUD-2, and LIN-5 in the presence of 10 µg/mL nocodazole. 100 × Real-Time. Scale bar: 10 µm.

**Movie S3.** Accumulation dynamics of dextrans (left) and HIS-58 (right). The molecular weights of dextran have been depicted in the movie. 100 × Real-Time.

**Movie S4.** Accumulation dynamics of NUD-2 and TBB-2 in *ran-1 (RNAi)* embryos. 200 × Real-Time. Scale bar: 10 µm.

**Movie S5.** Accumulation dynamics of NUD-2 fractions. 100 × Real-Time. Scale bars: 10 µm.
